# Hypoxia induces rapid changes to histone methylation reprogramming chromatin for the cellular response

**DOI:** 10.1101/513069

**Authors:** Michael Batie, Julianty Frost, Mark Frost, James W. Wilson, Pieta Schofield, Sonia Rocha

## Abstract

Molecular dioxygenases include JmjC-containing histone demethylases and PHD enzymes, but only PHDs are considered to be molecular oxygen sensors in cells. Although, it is known that hypoxia can alter chromatin, whether this is a direct effect on histone demethylases or due to hypoxia induced HIF-dependent transcriptional changes is not known. Here, we report that hypoxia induces a rapid and HIF-independent alteration to a variety of histone methylation marks. Genomic locations of H3K4me3 and H3K36me3 following short hypoxia predict the hypoxia gene signature observed several hours later in cells. We show that KDM5A inactivation mimics hypoxic changes to H3K4me3 in its targets and is required for the cellular response to hypoxia. Our results demonstrate a direct link between oxygen sensing and chromatin changes via KDM inhibition.

**One Sentence Summary:** Rapid oxygen sensing by chromatin

## Main Text

Hypoxia, or reduced oxygen availability, is an important stimulus for the development of multicellular organisms (*1*). In addition, hypoxia plays an important role in the pathology of numerous human diseases, contributing to treatment resistance and disease progression (*2*, *3*). At the molecular level, hypoxia activates a profound transcriptional programme in cells, with the ultimate aim of adaptation and survival at a lower oxygen level (*4*, *5*). To this end, the family of transcription factors, known as hypoxia inducible factors (HIFs), coordinate the majority of the transcriptional changes, with additional input from a multitude of additional and important transcription factors such as p53, NF-κB, Myc, and Notch (*4*, *6*). HIFs are heterodimeric transcription factors composed of HIF1β(oxygen insensitive) and HIF α(oxygen labile) (*7*). HIF α is encoded by three separate genes, HIF 1α, HIF 2α and HIF 3α, and their function in each tissue can be, at times, quite different (*7*). Hypoxia activation of HIFs is mediated via the inhibition of dioxygenases, most prominently prolyl-hydroxylases (PHDs) and Factor Inhibiting HIF (FIH) (*8*). These enzymes require molecular oxygen as a co-factor, as well as iron and 2-oxoglutarate, to conduct the hydroxylation reaction in HIF αproteins. Hydroxylation of prolines in HIF αby PHDs, creates a high affinity binding site for the tumour suppressor von Hippel Lindau (VHL), which forms part of a E3-ubiquitin ligase, promoting proteasome dependent and independent degradation of HIF α(*8*).

Analysis of chromatin marks, under conditions of hypoxia, has become an important aspect due to the discovery of a class of dioxygenases that can reverse histone methylation. These enzymes possess a Jumonji C domain (JmjC), which like PHDs and FIH require oxygen iron and 2-oxoglutarate to perform their enzymatic activity (*9*, *10*). As such, there is the potential for the activity of these enzymes to change in hypoxia, allowing specific areas of the genome to change in structure to allow transcriptional changes to occur. Interestingly, and much like PHD2 and PHD3, several of the JmjC containing enzymes have been shown to be direct HIF 1α targets (*10*), allowing for adaptation to occur at the chromatin level. Oxygen dependency for very few of the JmjC enzymes has been measured, and for these, the Km for oxygen is very similar to PHD enzymes *in vitro* (*11*, *12*), further suggesting that JmjC enzymes can act as molecular oxygen sensors in the cell. However, most of this information is missing for the vast majority of JmjC-histone demethylases. Interestingly, a number of studies have shown that histone methylation marks are increased following severe and prolonged hypoxia in a multitude of cell types (*10*, *13*, *14*).

Given that JmjC histone demethylases have the potential of being molecular oxygen sensors in the cell, we have investigated histone methylation changes under hypoxia at early stages of the hypoxia response. HeLa cells as well as the human fibroblast cell line HFF were exposed to hypoxia for a time course analysis. Using immunoblot analysis, it was possible to observe that hypoxia induced a rapid and robust increase in the levels of a variety of histone methylation marks on histone H3 (Fig. 1A), with no change in the levels of total histone H3 or actin in either cellular background. Interestingly, many of the changes observed in H3 methylation preceded robust stabilization of HIF 1α, occurring from 30 minutes following hypoxia exposure (Fig. 1A). Furthermore, increases in histone methylation were also seen after longer periods of hypoxia exposure (Fig. 1B). We also analysed if other HIF 1α stabilizing agents, such as DMOG (a known PHD inhibitor) and an iron chelator (DFX) could induce similar changes to histone H3 methylation marks (Fig. 1C-D). This analysis revealed a similar level and kinetics of histone H3 methylation increase as to that observed with hypoxia, suggesting that all these stimuli are working via a similar mechanism.

**Fig. 1.**
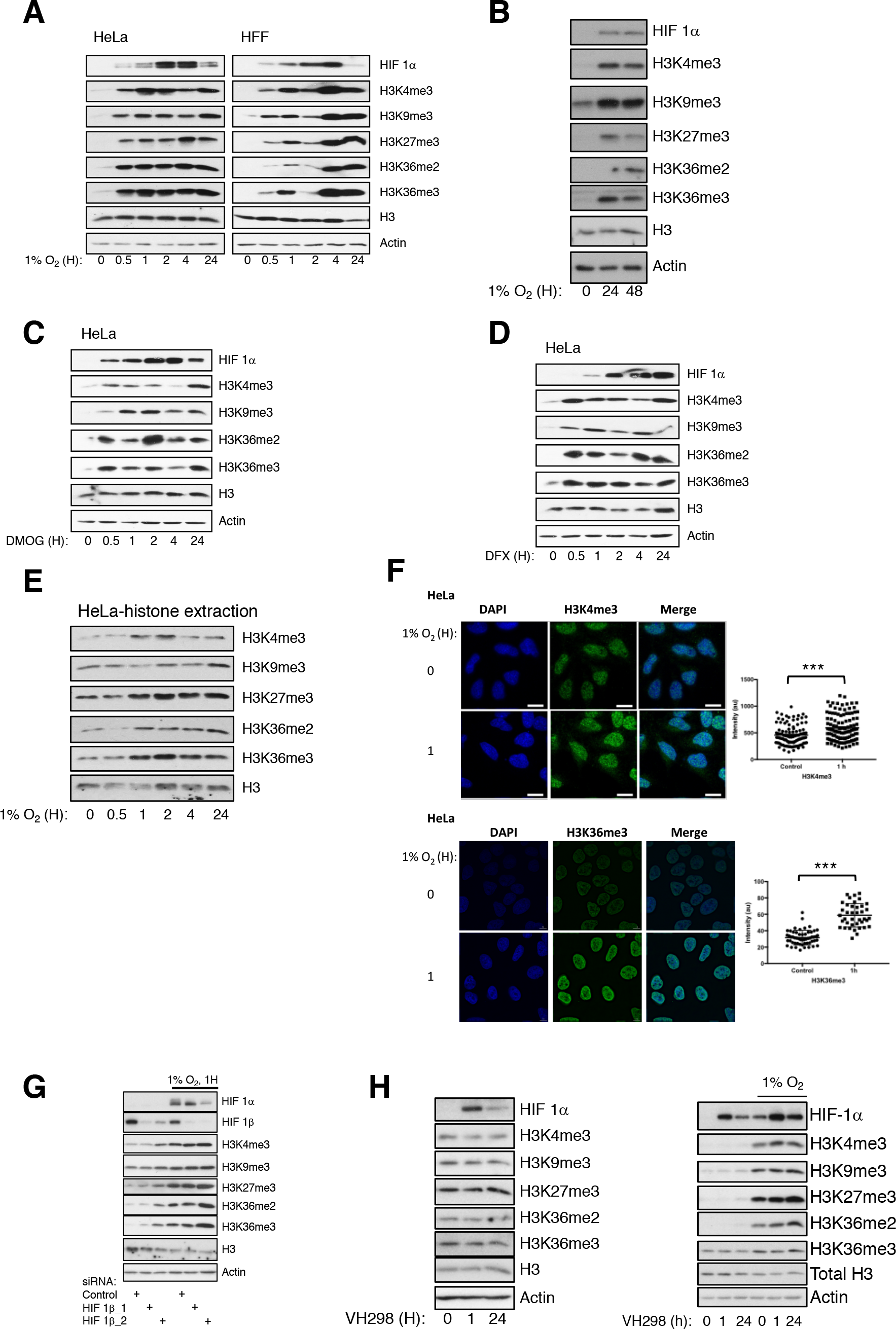
Hypoxia increases histone methylation independent of HIF. Immunoblot analysis of histone methylation and HIF 1α in HeLa and HFF cells exposed to 1% O_2_ for the indicated time points (**A**). Immunoblot analysis of histone methylation and HIF 1α in HeLa cells exposed to 1% O_2_ for the indicated time points (**B**). Immunoblot analysis of histone methylation and HIF 1α in HeLa cells treated with DMOG (**C**) and DFX (**D**) for the indicated time points. Immunoblot analysis of histone methylation on purified histone extracts from HeLa cells exposed to 1% O_2_ for the indicated time points (**E**). Immunofluorescence analysis histone methylation in HeLa cells exposed to 1% O_2_ for 1 hour (**F**) Graph depicts individual cell measurements, mean and SEM from a minimum of three independent experiments. ANOVA analysis determined statistical significance ***p< 0.001 between conditions. Immunoblot analysis of histone methylation, and HIF 1αand HIF 1βin HeLa cells exposed or not to 1 hour of 1% O_2_ following siRNA depletion of HIF 1β(**G**). Immunoblot analysis of histone methylation and HIF 1αin HeLa cells treated with VH298 for the indicated time points, with or without exposure to 24 hours to 1% O_2_ (**H**).

Prolonged hypoxia has also been associated with changes in reactive oxygen species (ROS) (*15*). We thus analysed if short-term hypoxia induced ROS in our cell system (Fig. S1A), and if using a ROS scavenger such as N-Acetyl-Cysteine (NAC) would alter histone methylation pattern in cells (Fig. S1B). Although we could detect a small increase in ROS after 1 hour 1% O_2_ (Fig. S1A), pre-treatment with NAC did not reduce hypoxia induced increases in histone methylation (Fig. S1B). In addition, when cells were treated with high levels of H_2_O_2_, to mimic ROS production, we could not detect changes in histone methylation marks following 1 hour but increases were visible following 24 hours (Fig. S1C). These data suggest that ROS is not involved in the increase of histone methylation marks observed at 1 hour hypoxia, but could contribute to increases in histone methylation following prolonged hypoxia.

Another known associated factor with hypoxia exposure is changes in metabolism, and hence increased levels of certain metabolites (*16*). Of particular relevance to histone demethylases are changes in 2-oxoglutarate related metabolites such as succinate and fumarate and also the oncometabolite 2-hydroxyglutarate (2-HG) (*16*, *17*). In fact, recent studies have measured the affinity of certain histone demethylases for such types of metabolites but also measured cells responses to treatment with metabolites and oncometabolites (*17*). However, these studies relied on prolonged exposure to these, sometimes up to 72 hours of treatment (*17*). To determine if short hypoxia induced changes in histone methylation were mimicked by changes in metabolites or oncometabolites, we treated HeLa cells with succinate, fumarate or cell permeable 2-HG (Fig. S2A-C). While we could not detect any changes in the histone methylation marks we analysed following treatment with succinate (Fig. S2A), fumarate could induce some increases in histone methylation (Fig. S2B). Treatment with 5 mM 2-HG resulted in increases in methylation marks in histone H3 only following prolonged treatment, but not after 1 hour (Fig. S2C). However, this was only seen for HeLa cells, while in HFF cells, 2-HG treatment, even after prolonged exposure, only altered H3K4me3 and H3K27me3 (Fig. S2C). Finally, we measured levels of 2-HG in cells exposed to 1 hour or 24 hours of hypoxia (Fig. S2D). As a positive control, we measured 2-HG levels in HT1080 cells that possess a mutation in IDH1, which results in high levels of this oncometabolite (*18*). This analysis revealed that in HeLa or HFF cells, hypoxia treatment, at the time points we analysed, did not result in any significant change to 2-HG levels, suggesting that this oncometabolite is not involved in the mechanism controlling the observed changes to the histone marks.

Since our lysis conditions extract both soluble and most of the chromatin bound proteins, we next determined if hypoxia was leading to increased histone H3 methylation in chromatin bound only histones. To that end, we performed acid extraction of histone and re-analysed H3 methylation marks by immunoblot (Fig. 1E, Fig. S3). It was possible to determine that following 1 hour of hypoxia exposure, there were visible increases in several H3 methylation marks (Fig. 1E, Fig. S3). This indicates that indeed chromatin bound H3 possesses higher levels of methylation following hypoxia. Finally, we supplemented our immunoblot analysis with quantitative immunofluorescence (Fig. 1F). Here, short term exposure to hypoxia resulted in a significant increase to the levels of H3K36me3 and H3K4me3.

Although these results suggest that HIF is not involved in the increases in histone methylation in hypoxia, it is important to demonstrate this formally. To this end, we depleted cells of HIF 1β using two separate siRNAs and analysed levels of the histone methylation on histone H3 following 1 hour of hypoxia exposure (Fig. 1G). Surprisingly, depletion of HIF 1β resulted in even higher levels of histone methylation marks in hypoxia, demonstrating the HIF is not required for the changes observed in hypoxia. Increased levels of histone marks in the absence of HIF 1α could be due to reduced levels of several JmjC histone demethylases since these are known to be direct targets of the HIF 1 complex (*10*, *19*). To further investigate the involvement of HIF in our mechanism, we used the selective chemical probe VH298 (*20*), which inhibits VHL without altering PHD or other dioxygenases functions in cells (*20*). Here, despite robust HIF 1α stabilization, no changes in histone methylation marks were observed in both normoxia and hypoxia (Fig. 1H). Whereas, using cells constitutively expressing active HIF 1α and/or HIF 2α, hypoxia was able to induce increases in histone methylation marks (Fig. S4A-C). These data indicate that hypoxia, and other dioxygenase inhibitors induce rapid changes to several histone methylation marks but HIF is not required for this response.

The analysis on H3 methylation suggests that hypoxia induces global changes to methylation marks. However, two possible consequences could result from these changes. One would be a stochastic distribution of the increased H3 methylation across the genome or alternatively, loci specific and gene expression directed changes would occur following rapid exposure to hypoxia. To determine which of these two options is occurring and in an unbiased manner, we performed Chromatin Immunoprecipitation (ChIP) followed by next generation sequencing for H3K4me3 and H3K36me3 marks in normoxia and following 1 hour exposure to 1% O_2_ in HeLa cells.

Analysis of H3K4me3 data revealed that in all 3 biological replicates, we could identify over 14,000 peaks in both conditions, corresponding to 11,000 genes (Fig. S5A-B, Dataset S1). Of all the peaks identified for both treatment conditions across 3 biological replicates, over 12,000 were shared between normoxia and hypoxia (Fig. 2A). In addition, when compared to a dataset produced by the ENCODE consortium, this covered 92% of all peaks in normoxia (Fig. S5C). The hypoxia dataset was covered by 91% of the dataset in ENCODE (Fig. S5C). This indicated that our dataset is of high quality, comparable to the standard of ENCODE. Using stringent analysis conditions, we could determine that hypoxia exposure led to 164 peaks upregulated and 455 peaks that were reduced (Fig. 2B, Dataset S1). Analysis of genomic location of all H3K4me3 peaks present in all three biological replicates reveal similar distributions with 95% of all peaks locating to promoter regions both in normoxia and hypoxia, within 3000bp from the transcription start site (TSS) (Fig. S5D). H3K4me3 increased peaks were not only located within promoter regions but had also increased localization in gene bodies and intergenic regions (Fig. S5E). On the other hand, hypoxia reduced peaks were almost exclusively located to promoters (Fig. S5E). To understand if the intergenic or gene body location of the increased H3K4me3 peaks had some functional significance, we determined if these sites were associated with enhancer markers. From our 25 peaks, identified in these non-promoter locations, 80% maps to DNAse hypersensitive sites and 60% to sites of H3K27Ac. Furthermore, 25% were also identified in the Fantom5 dataset of predicted enhancers (Fig. S5F). This suggests that increased H3K4me3, at these intergenic regions, could be controlling expression of associated genes. Given that none of these genes linked to enhancers, identified in the Fantom5 analysis, have been shown to be hypoxia responsive, we performed qPCR analysis in HeLa cells following several time points of exposure to 1% O_2_ (Fig. S6A). This revealed that CBLB, CGAS and EEF1A1 are hypoxia inducible after 24 hours in HeLa cells, supporting the hypothesis that H3K4me3 increases at enhancer regions associated with these genes could control their expression.

**Fig. 2.**
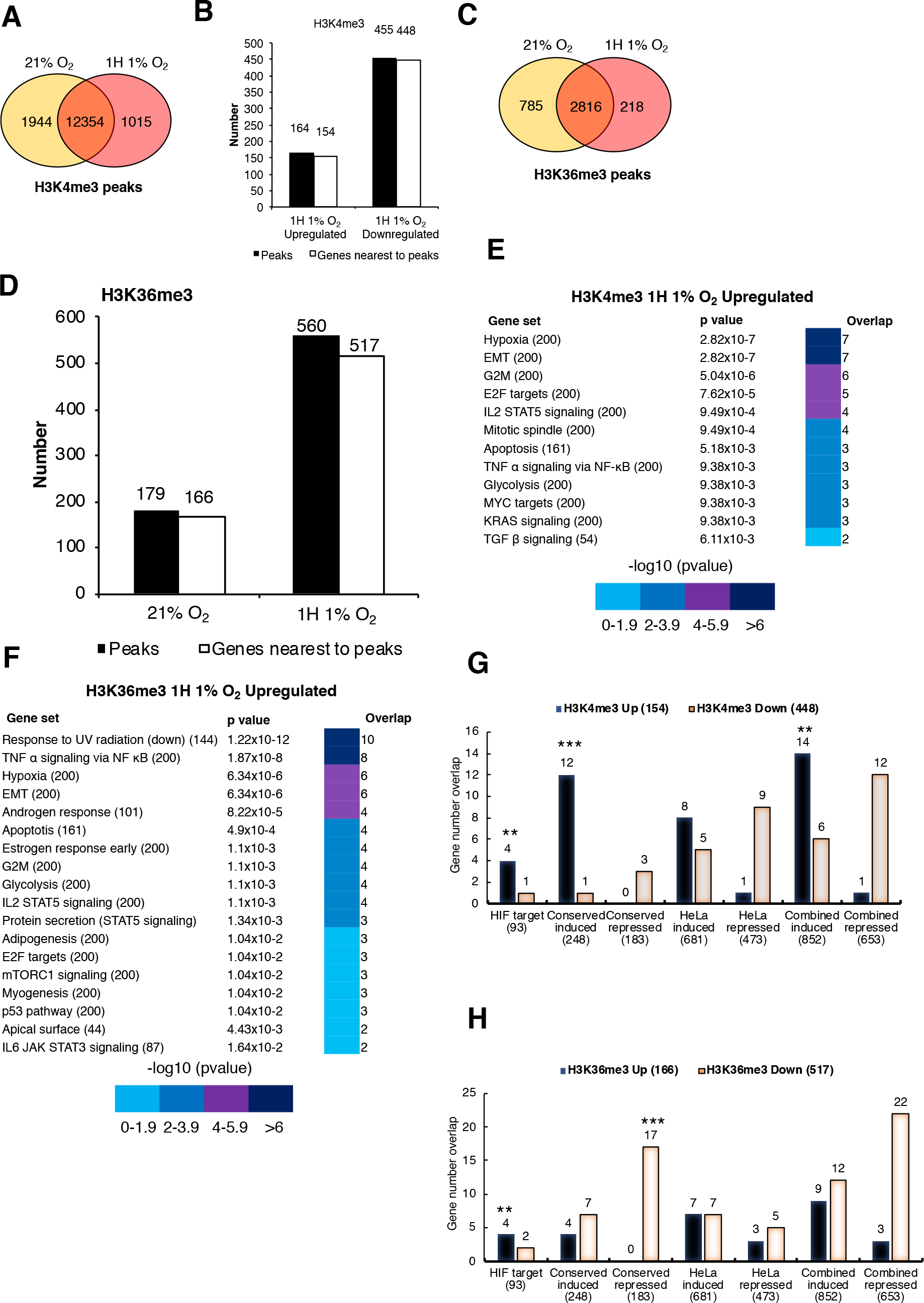
Hypoxia induces specific changes in H3K4me3 and H3K36me3 with marks increasing at hypoxic gene signatures. ChIP-Seq analysis of H3K4me3 and H3K36me3 in HeLa cells at 21% O_2_ (normoxia) or exposed to 1% O_2_ for 1 hour (hypoxia). Overlap of high stringency (3/3 replicates) H3K4me3 (**A**) and H3K36me3 (**C**) ChIP-seq peaks in HeLa cells at normoxia (21% O_2_) or exposed to hypoxia (1H of 1% O2). Differential regulation of H3K4me3 (**B**) and H3K36me3 (**D**) ChIP-seq peaks in HeLa cells exposed to hypoxia compared to normoxia. Gene group association analysis showing significant enrichment of gene set signatures (MsigDB) for H3K4me3 (**E**) and H3K36me3 (**F**) hypoxia upregulated ChIP-Seq peak genes. Hypergeometric distribution analysis determined statistical significance. Gene group association analysis of differentially regulated H3K4me3 (**G**) and H3K36me3 (**H**) ChIP-Seq peaks in HeLa cells exposed to hypoxia (1H, 1% O_2_) compared to normoxia, with HIF 1αtarget genes (HIF target), hypoxia induced genes, hypoxia repressed genes, hypoxia induced and repressed genes in HeLa cells, and the non-redundant combination of HIF 1αtarget and hypoxia induced genes (HIF 1αtarget + hypoxia induced). p values were determined by Fisher exact t test. **p< 0.01, ***p< 0.001.

Given the nature of H3K36me3 mark distribution, we performed a high stringency peak calling analysis. This gave rise to the identification of 3829 peaks called in normoxia, and 3125 peaks called in 1% O_2_, seen across all three biological replicates, corresponding to ~3,000 genes (Fig. S7A-B, Dataset S1). Overall, 2816 H3K36me3 peaks were found in normoxia and hypoxia (Fig. 2C). Comparison with ENCODE data for this mark again indicated the quality of our dataset, with normoxia and hypoxia covering 92 % of peaks present in the ENCODE dataset (Fig. S7C). Hypoxia led to 179 peaks being upregulated and 560 peaks being downregulated (Fig. 2D, Dataset S1). H3K36me3 located primarily across the gene bodies (Fig. S7D). Hypoxia led to a reduction in the percentage of peaks identified for this mark, but its location was unaltered (Fig. S7E).

To understand the relevance of the changes in H3K4me3 and H3K36me3 we had identified after such a short period of hypoxia, we performed pathway association analysis using the molecular signatures database (MSigDB) (*21*, *22*). Genes found to have upregulated peaks of H3K4me3 had a significant representation of hypoxia signaling and epithelial to mesenchemical transition signatures (Fig. 2E). On the other hand, genes with reduced peaks of H3K4me3, were mostly associated with cell division and oxidative phosphorylation (Fig. S8A). Analysis of the H3K36me3 datasets revealed that genes with increased H3K36me3 peaks were associated not only with hypoxia but also NF-κB signaling (Fig. 2F). Although ‘Response to UV’ (downregulated) was the signature mostly represented in this dataset, a similar signature was also found associated with the genes with reduced peaks for H3K36me3 (Fig. S8B). Genes with reduced H3K36me3 peaks were also associated with cell division and oxidative phosphorylation, suggesting that these molecular pathways are represented in the acute response to hypoxia (Fig. S8B).

Given the interesting results that 1 hour of hypoxia exposure changes histone mark deposition at genes associated with the hypoxia response, we performed integrative analysis of our histone marks datasets, with known HIF targets and hypoxia inducible/repressed gene sets (Fig. 2G-H). Overlap analysis was performed on 93 validated HIF 1 target genes (adapted from (*23*)), hypoxia induced and repressed genes across several cell types (*24*) (Fig. 2G, Dataset S1). H3K4me3 hypoxia upregulated peaks were significantly correlated with validated HIF targets, cell type conserved hypoxia induced genes and the non-redundant combination of validated HIF targets and cell type conserved hypoxia induced genes (Fig. 2G, Dataset S1). H3K4me3 hypoxia downregulated peak genes showed no significant correlation with HIF validated target genes, hypoxia induced or hypoxia repressed gene lists (Fig. 2G). We also compared the H3K4me3 ChIP-seq dataset with our own RNA-seq analysis in HeLa cells, following 16 hours exposure to 1% O_2_ (Fig. 2G, Dataset S1). This analysis revealed some overlap with hypoxia induced genes, but also contained genes that were identified as having a reduced level of H3K4me3 but were still induced in HeLa cells at 16 hours. This result was not unexpected given the dynamic nature of gene expression in hypoxia, with certain genes being activated at early times but repressed at later stages, when adaptation is starting to occur. Finally, H3K4me3 hypoxia upregulated peaks were significantly enriched in the non-redundant combination of validated HIF target genes, cell type conserved hypoxia induced genes and HeLa hypoxia induced genes. Taken together these results suggest that H3K4me methylation is associated with a core response to hypoxia, since most significant overlap occurs in a cell independent and conserved set of genes responsive to this stress.

H3K36me3 upregulated peak genes also displayed positive correlation with HIF validated target genes (Fig. 2H, Dataset S1). All of these peak changes were located at gene bodies (Dataset S1). No gene overlap was found between H3K36me3 hypoxia upregulated peak genes and cell type conserved hypoxia downregulated genes (Fig. 2H). However, hypoxia repressed genes were significantly enriched for H3K36me3 hypoxia downregulated peak genes (Dataset S1).

For validation, we focused on the H3K4me3 genes, selecting some known HIF genes but also genes that are hypoxia-inducible independently of known HIF regulation (Dataset S1). We also included a very well-known HIF-dependent and hypoxia inducible gene, not identified in our stringent analysis, *CA9* (Fig. S9A). ChIP-qPCR analysis revealed a significant increase in H3K4me3 present at all the predicted genes, with further increases being observed at 24 hours of hypoxia (Fig. 3A, Fig. S9B). Interestingly, we also observed significant increases in the mark present at the *ACTB* gene, which we used as a control, following 24 hours of hypoxia. As negative controls we used BAP1 and KDM2B genes, BAP1 was identified as a down peak while no change was detected in KDM2B in the H3K4me3 analysis. As predicted no change was observed in the levels of H3K4me3 present at the genomic locations analysed following any of the hypoxia time points (Fig. 3A, Fig. S9B)

**Fig. 3.**
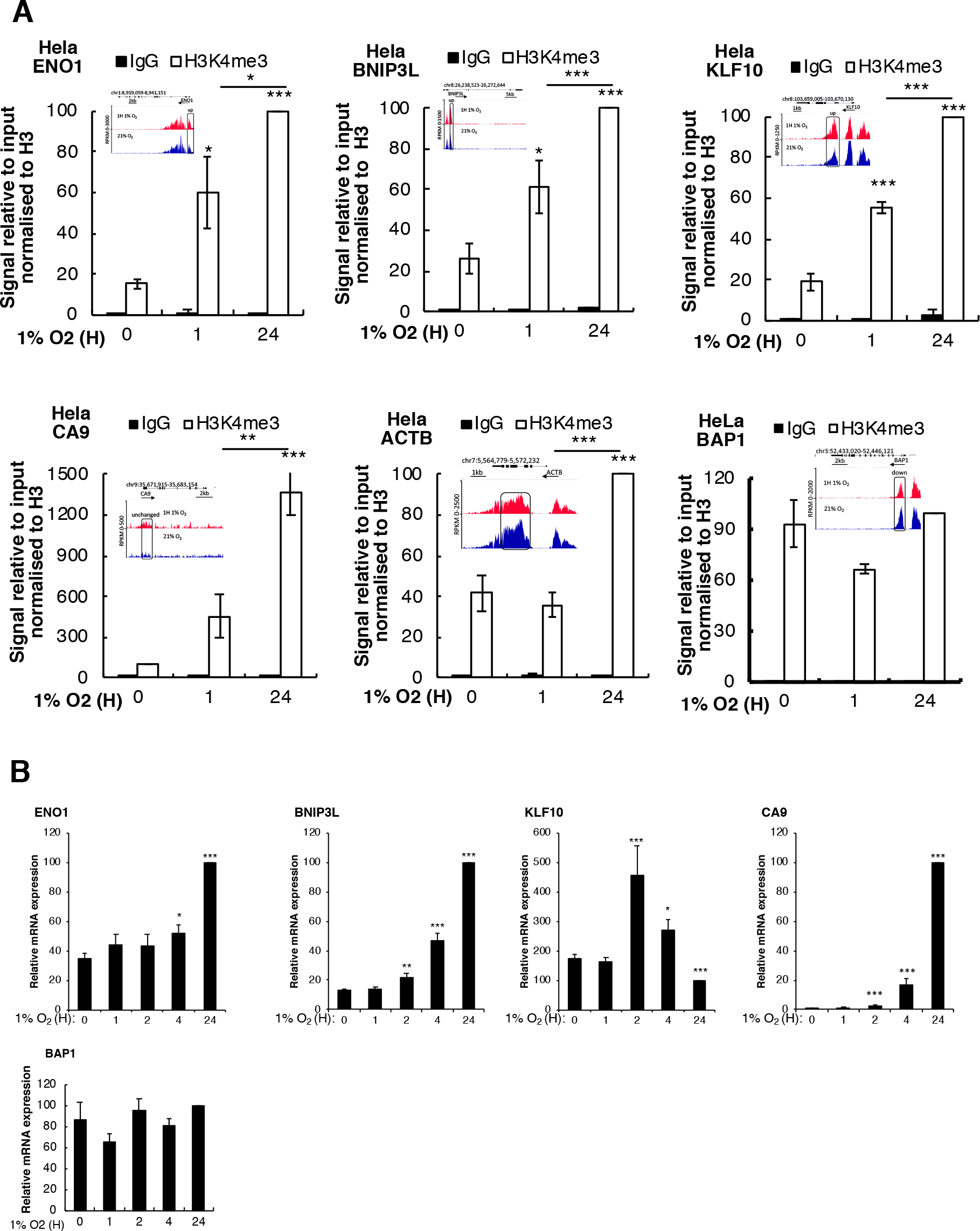
H3K4me3 levels increase in hypoxia at a subset of hypoxia inducible genes. ChIP qPCR analysis of H3K4me3 at promoters of indicated the genes in HeLa cells exposed to 1% O_2_ for the indicated time points (**A**). Coverage tracks for RPM normalised H3K4me3 ChIP-seq read counts in HeLa cells exposed to normoxia (N) or 1 hour of 1% O_2_ (H) with hypoxia upregulated peaks highlighted, are inserted into graphs. Data represent the mean and SEM of at least 3 independent experiments with p values determined by Student’s t test. (**B**) qPCR analysis of mRNA expression levels for the indicated genes in HeLa cells exposed to 1% O_2_ for the indicated time points. Data represent the mean and SEM of at least 3 independent experiments with p values determined by Student’s t test. *p< 0.05, **p< 0.01, ***p< 0.001.

Since H3K4me3 is associated with active transcription, we next determined if the genes selected for validation of our sequencing results were being activated very early in hypoxia. To this end, we analysed mRNA levels for all the genes we tested using qPCR (Fig. 3B, Fig. S10A). This analysis demonstrated that following 1 hour of hypoxia, none of the investigated genes increased above the normoxia control. *BNIP3L*, *KLF10, CA9* and *STAG2* mRNA levels were significantly increased following 2 hours of hypoxia, while *ENO1* and *LOX* mRNA levels were only increased following prolonged hypoxia exposures. These results indicate that elevated H3K4me3, following 1 hour of hypoxia, is not a result of increased gene transcription at the sites studied.

Given the rapid nature of the changes in H3K4me3 levels observed at these genes, we hypothesized that this was due to JmjC histone demethylase inhibition. This mark is predicted to be erased by a variety of these enzymes, namely, KDM2B and the KDM5A-D family. To investigate if any or all of the enzymes were controlling global levels of this mark in HeLa cells, we depleted each of them using siRNA and analysed the levels of H3K4me3 by immunoblot (Fig. 4A, Fig. S10B-C). Visible increases in H3K4me3 could be seen in cells depleted of KDM2B, KDM5A, KDM5B and KDM5C (Fig. 4A). However, no effect was seen with KDM5D, which is not expressed in HeLa cells (Fig. 4A). We also analysed mRNA levels of the genes we identified as having increased levels of H3K4me3 at their promoters in hypoxia (Fig. 4B, Fig. S10D). This revealed that KDM5A has the strongest effect, inducing increases in mRNA levels of BNIP3L, KLF10 and LOX. STAG2 and ENO1 mRNA levels were only increased significantly by one of the siRNA oligonucleotides used against KDM5A (Fig. 4B, Fig. S10D). These changes were mirrored in the levels of H3K4me3 present at promoters of each of the genes tested, with significant increases detected in KLF10, BNIP3L, STAG2 and LOX, when cells were depleted of KDM5A but not in our negative controls BAP1 and KDM2B (Fig. 4C, Fig. S10E). These results indicate that KDM5A is directly involved in the regulation of H3K4me3 at the genes we identified in the ChIP-sequencing in the early response to hypoxia. KDM5A involvement in hypoxia had previously been demonstrated following prolonged exposures times in other systems (*25*).

**Fig. 4.**
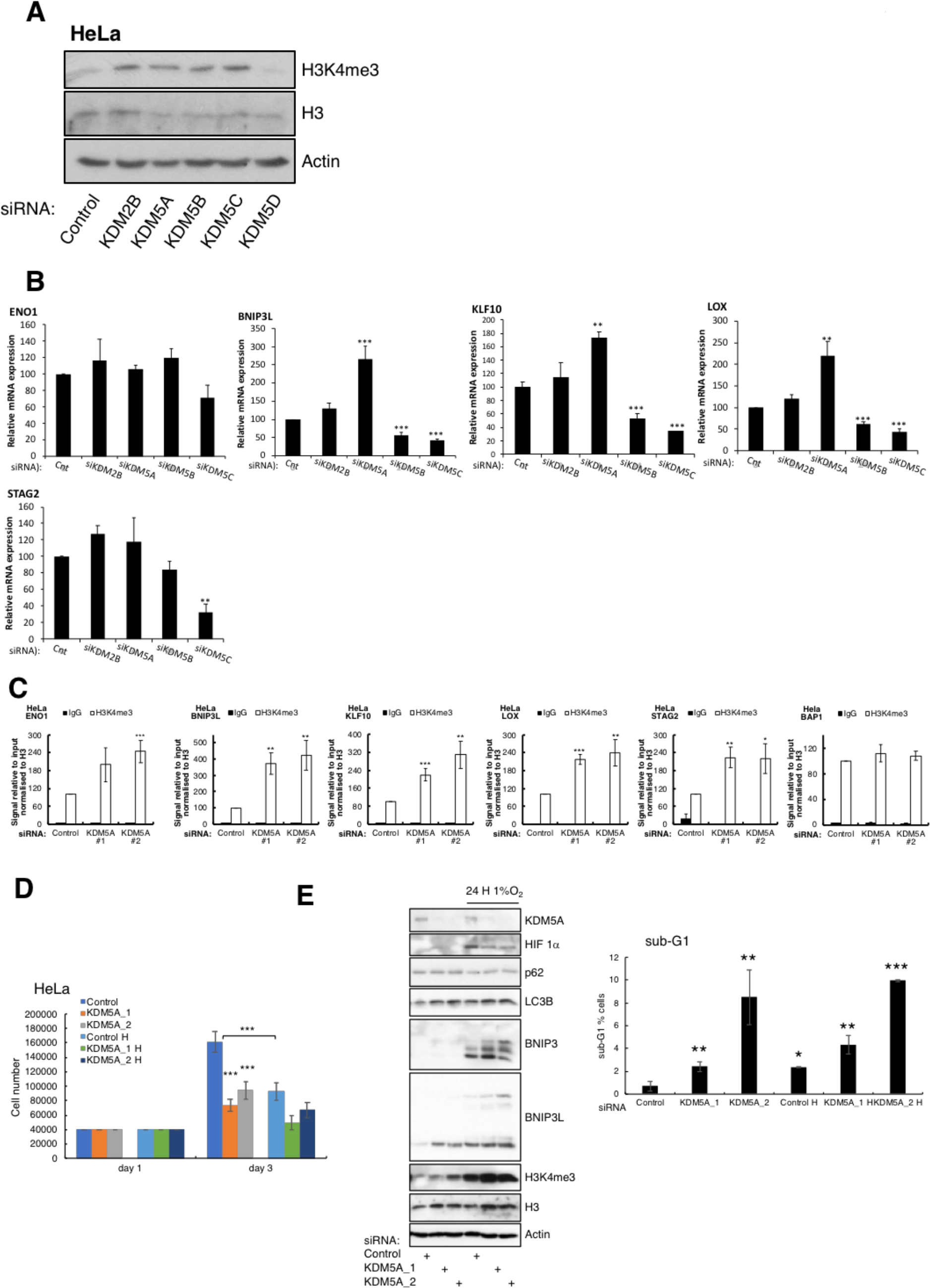
KDM5A regulates promoter methylation and gene expression at a subset hypoxia inducible genes and controls cells proliferation and autophagy. Immunoblot analysis of H3K4me3 and H3 levels in HeLa cells siRNA depleted of KDM5A, KDM5B, KDM5C and KDM5D (**A**). qPCR analysis of mRNA expression levels for the indicated genes in HeLa cells siRNA depleted of KDM5A, KDM5B and KDM5C (**B**). Data represent the mean and SEM of at least 3 independent experiments with p values determined by Student’s t test. ChIP qPCR analysis of H3K4me3 at promoters of indicated the genes in HeLa cells siRNA depleted of KDM5A (**C**). Data represent the mean and SEM of at least 3 independent experiments with p values determined by Student’s t test. Cell proliferation analysis in HeLa cells depleted of KDM5A in normoxia and hypoxia (**D**). Analysis of autophagy markers and BH3 protein BNIP3 and BNIP3L by immunoblot in cells depleted of KDM5A and exposed to hypoxia and sub-G1 analysis using Flow cytometry (**E**). Data represent the mean and SEM of at least 3 independent experiments with p values determined by Student’s t test. *p< 0.05, **p< 0.01, ***p< 0.001.

To determine if indeed KDM5A has a role in altering cellular responses in hypoxia and normoxia, we performed functional assays modulating KDM5A levels in cells. We could demonstrate that KDM5A depletion resulted in reduced proliferation and colony formation in normoxia but also hypoxia (Fig. 4D, Fig. S10F). Furthermore, KDM5A depletion increased the levels of Autophagy/Apoptosis BH3 containing proteins both in normoxia and hypoxia (Fig. 4E). As predicted by our ChIP-seq validation and mRNA analysis, KDM5A depletion is sufficient to induce BNIP3L protein, an important BH3 containing protein with roles in autophagy and apoptosis (*26*, *27*). These results demonstrate that KDM5A is sensitive to oxygen levels in cells and modules the cell’s transcription and functional response.

Our results indicate that chromatin can sense oxygen at the cellular level, via JmjC domain containing enzymes, in a similar manner to HIF 1α via the PHD enzymes. We thus investigated the dynamics of histone methylation marks in hypoxia. A feature of oxygen sensing by PHDs is the rapid re-instatement of basal HIF 1α levels, upon re-oxygenation (*28*). We thus analysed how rapidly histone methylation marks retuned to normoxic levels upon restoration of normal oxygen tensions (Fig. 5A). As expected, HIF 1α levels quickly returned to basal or lower levels when cells were exposed to 21% O_2_ following 24 hours exposure to 1% O_2_ (Fig, 5A). Interestingly, almost all histone marks were reverted as quickly as 1 hour following re-oxygenation, indicating again a strong sensitivity to O_2_ tensions at this level (Fig. 5A). To further delineate the O_2_ sensitivity of histone methylation marks to O_2_, we repeated our analysis in cells exposed to different levels of O_2_ and compared this to HIF 1α stabilization pattern in the cell (Fig. 5B). This revealed that the histone methylation marks we analysed are increased at the same O_2_ concentration as HIF 1α, indicating that some of the JmjC containing enzymes have similar O_2_ requirements as the PHD enzymes (Fig. 5B). This is in agreement with the reported Km measurement reported for of these enzymes, KDM4A and KDM4E (*11*, *12*).

**Fig. 5.**
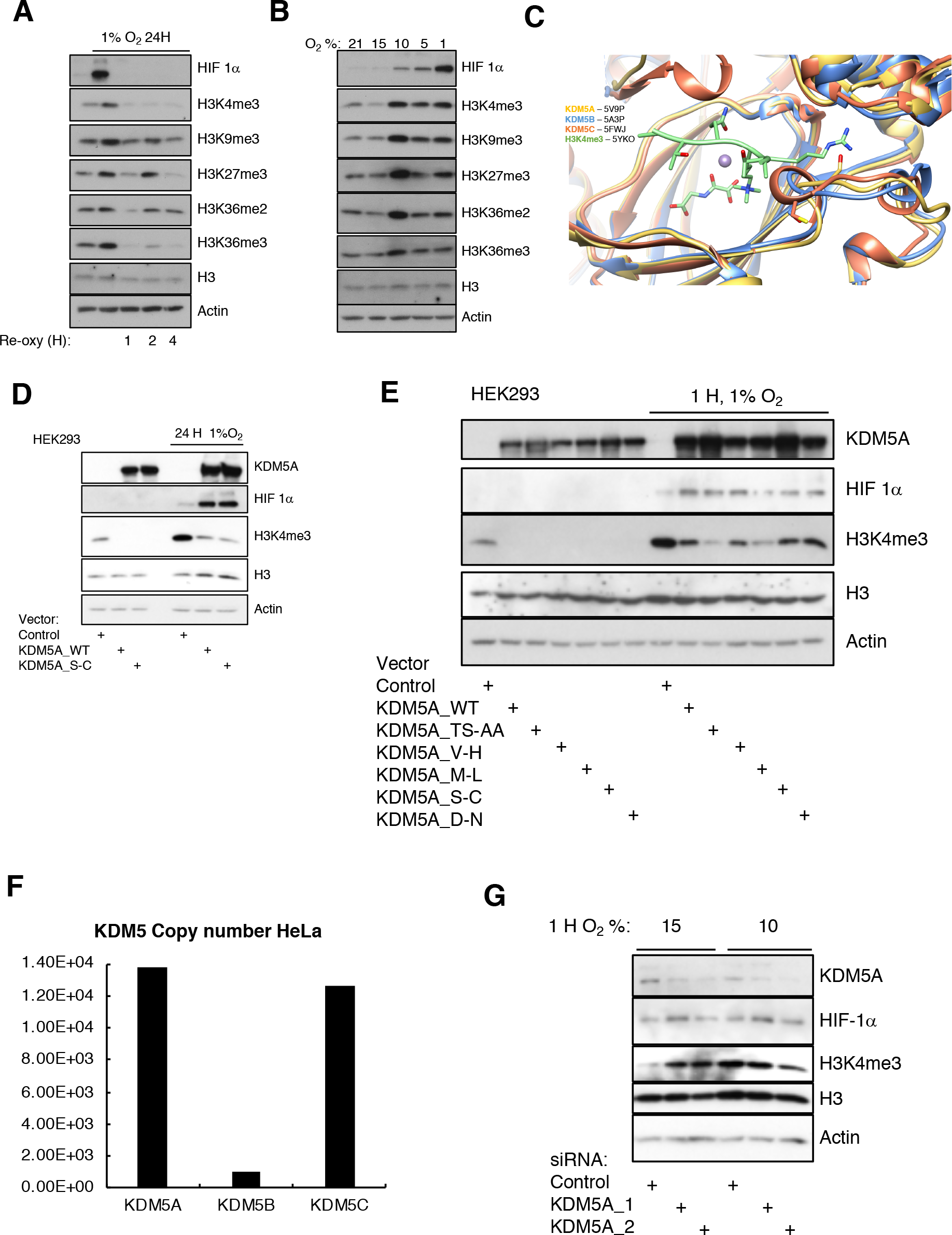
Oxygen sensitivity of cellular histone methylation marks. Immunoblot analysis of histone methylation in HeLa cells exposed or not to 24 hours of 1% O_2_ followed by 1,2 and 4 hours of reoxgenation at normoxia (21% O_2_) (**A**). Immunoblot analysis of histone methylation in HeLa cells exposed or not to 1 hour of 15%, 10%, 5% and 1% O_2_ (**B**). Structural alignment of KDM5A (yellow, PDB 5V9P), KDM5B (blue, PDB 5A3P), and KDM5C (orange, PDB 5FWJ) catalytic pocket with the H3K4me3 peptide substrate (green, PDB 5YKO) (**C**). Immunoblot analysis of H3K4me3 levels in the presence of overexpressed KDM5A wildtype or Serine 464 to Cysteine mutant in HEK293 cells. Where indicated cells were also exposed to 24 hours of 1% O_2_ (**D**). Immunoblot analysis of H3K4me3 levels in the presence of overexpressed KDM5A wildtype or the indicated KDM5A mutants in HEK293 cells. Where indicated cells were also exposed to 24 hours of 1% O_2_ (**E**). Quantitative proteomic analysis for KDM5 family members in HeLa cells (**F**). Immunoblot analysis of histone methylation and HIF 1αlevels in HeLa cells siRNA depleted of KDM5A and exposed or not to 15% and 10% O_2_ for 1 hour (**G**).

Given that our results indicated that KDM5A is the more likely KMD5 family member to be mediating the O_2_ sensing in HeLa cells, we investigated if there were any sequence or structural differences between KDM5A, KDM5B and KDM5C that could explain these observations. Sequence alignment revealed a high level of similarity between all three enzymes (Fig. S11A). Closer examination of the KDM5 family active site using publicly available structures, revealed very similar features (Fig. 5C). The only visible difference is on Serine 464 in KDM5A and KDM5C (position 495), which is changed to a Cysteine in KDM5B (position 480), this creates a bigger onward interface in KDM5A than in the other KDM5 family members (Fig. 5C). We thus created this point mutation in KDM5A and assessed its response in normoxia and hypoxia (Fig. 5D). For this analysis we used HEK293 cells, which also demonstrably respond with similar kinetics of histone demethylase inhibition as HeLa cells (Fig. S12A). Overexpression of KDM5A resulted in robust demethylation in normoxia as expected (Fig. 5D). KDM5A was also able to demethylate H3K4me3 even at 1% O_2_, suggesting that increasing the levels of the enzyme can compensate for reduced level of co-activators. Interestingly, mutation of Serine 464 to Cysteine did not results in any discernable change to the activity of KDM5A, suggesting that this site is not responsible for the specificity we are proposing in response to hypoxia. We thus returned to the structural information available and created a series of mutants across different and divergent regions of KDM5A (Fig. S12B). These mutations were directed against key domains, where residues were different between KDM5 family members (Fig. S12B). We analysed the activity of these mutants in cells in normoxia and 1% O_2_. All mutants expressed well, and similar potent at demethylating H3K4me3 in normal oxygen conditions (Fig. 5E). As previously observed, mutation of S464 to Cysteine did not result in a visible activity change in either normoxia or hypoxia. Similar results were obtained for mutations in the ARID domain or PHD3 domain (Fig. 5E, Fig. S12B). Interestingly, mutations in the JmjN (T30 and S34 to A) and PHD1 (M297L) domains resulted in higher demethylase activity even at 1% oxygen. Given that these domains do not coordinate coactivators such as oxygen or 2-oxoglutarate, it is likely that they increase affinity for histone tail (PHD1 domain) and increase activity by potentiating dimerization (ARID domain), as was recently discovered for the KDM4 family (*29*).

The mutational analysis of KDM5A, making it more similar to KDM5B, still could not answer how the specificity of oxygen dependent responses is achieved in cells. We hypothesized that this could be due to levels of the respective enzymes in cells. HIF 1α stabilization in cells is only transient given the negative feedback loop mediated by PHD2 and PHD3. After prolonged hypoxia, upregulation of PHD2 or PHD3 in a HIF 1α dependent manner, results in reduced HIF 1α levels, despite low oxygen conditions. It is therefore possible that different levels of KDM family members can dictate specificity of a cell/tissue response to oxygen. Based on this hypothesis, we interrogated quantitative proteomic datasets available on the PRIDE database and collected the copy number information for several members of the KDM family in HeLa cells (*30*) (Fig. S13A-E). In HeLa cells, KDM5A is the most abundant KDM5 family member, followed closely by KDM5C, while KDM5B is much less abundant (Fig. 5F). Clear copy number differences amongst KDM family members is seen for all the KDMs detected in these proteomic analyses, with KDM2A, KDM3B, KDM4A, and KDM6A having highest expression levels in HeLa cells (Fig. S13A-E). Our overexpression analysis (Fig. 5D-E), already supports this possibility. However, we manipulated KDM5A levels with siRNA and exposed cells to 15% or 10% of O_2_ and analysed how this histone mark behaved (Fig. 5G). This analysis revealed that reduced levels of KDM5A, and hence activity, resulted in similar H3K4me3 levels in cells exposed to 15% or 10% O_2_, thus setting a new sensitivity threshold for oxygen in these cells (Fig. 5G). Although the O_2_ Km for KDM5A is currently unknown, our data suggest that this property alone is not sufficient to explain how different tissues respond to different O_2_ tensions. Rather, and based on the data presented here, levels of JmjC histone demethylases in combination with oxygen affinity, contribute to the overall sensitivity to O_2_ of a given cell, and this would promote specificity across different tissues. JmjC containing enzymes, including KDM5A, are evolutionary conserved from yeast to humans (*31–33*) suggesting that oxygen sensing by chromatin via these enzymes will contribute to a conserved response to hypoxia across species. Furthermore, it suggests that oxygen sensing by chromatin is, in phylogenetic evolutionary terms, older that oxygen regulated transcription factors such as HIFs.

## Acknowledgments

We thank Shaun Cottrill and Andrew Clark for technical assistance and Anne Hermann for help with Fluorometric measurements. We also thank the Centre for Cell Imaging and the accounts team in IIB at the University of Liverpool for help.

## Author Contributions

MB and SR conceived the experiments and wrote the manuscript. MB, JF, MF, JWW, and SR performed experiments. MB, JF, MF, PS, and SR analysed the data.

## Data and materials availability

All data is available in the main manuscript and supplementary material. ChIP-Seq and RNA-seq data has been uploaded to the NCBI GEO.

## Supplementary Materials

Materials and Methods

Figures S1-S13

Dataset S1

References (*34-57*)

## References and Notes

1. W. G. Kaelin, Jr., P. J. Ratcliffe, Oxygen sensing by metazoans: the central role of the HIF hydroxylase pathway. Mol Cell 30, 393–402 (2008).

2. T. G. Smith, P. A. Robbins, P. J. Ratcliffe, The human side of hypoxia-inducible factor. Br J Haematol 141, 325–334 (2008).

3. P. J. Pollard et al., Regulation of Jumonji-domain-containing histone demethylases by hypoxia-inducible factor (HIF)-1alpha. The Biochemical journal 416, 387–394 (2008).

4. N. S. Kenneth, S. Rocha, Regulation of gene expression by hypoxia. The Biochemical journal 414, 19–29 (2008).

5. M. Batie, L. Del Peso, S. Rocha, Hypoxia and Chromatin: A Focus on Transcriptional Repression Mechanisms. Biomedicines 6, (2018).

6. E. P. Cummins, C. T. Taylor, Hypoxia-responsive transcription factors. Pflugers Arch 450, 363–371 (2005).

7. S. Rocha, Gene regulation under low oxygen: holding your breath for transcription. Trends Biochem Sci 32, 389–397 (2007).

8. W. G. Kaelin, Proline hydroxylation and gene expression. Annu Rev Biochem 74, 115–128 (2005).

9. A. Melvin, S. Rocha, Chromatin as an oxygen sensor and active player in the hypoxia response. Cellular signalling 24, 35–43 (2012).

10. A. Shmakova, M. Batie, J. Druker, S. Rocha, Chromatin and oxygen sensing in the context of JmjC histone demethylases. The Biochemical journal 462, 385–395 (2014).

11. E. M. Sanchez-Fernandez et al., Investigations on the oxygen dependence of a 2-oxoglutarate histone demethylase. The Biochemical journal 449, 491–496 (2013).

12. R. L. Hancock, N. Masson, K. Dunne, E. Flashman, A. Kawamura, The Activity of JmjC Histone Lysine Demethylase KDM4A is Highly Sensitive to Oxygen Concentrations. ACS chemical biology 12, 1011–1019 (2017).

13. M. E. Adriaens et al., Quantitative analysis of ChIP-seq data uncovers dynamic and sustained H3K4me3 and H3K27me3 modulation in cancer cells under hypoxia. Epigenetics Chromatin 9, 48 (2016).

14. P. Prickaerts et al., Hypoxia increases genome-wide bivalent epigenetic marking by specific gain of H3K27me3. Epigenetics & chromatin 9, 46 (2016).

15. K. A. Smith, G. B. Waypa, P. T. Schumacker, Redox signaling during hypoxia in mammalian cells. Redox Biol 13, 228–234 (2017).

16. L. Tretter, A. Patocs, C. Chinopoulos, Succinate, an intermediate in metabolism, signal transduction, ROS, hypoxia, and tumorigenesis. Biochim Biophys Acta 1857, 1086–1101 (2016).

17. T. Laukka, M. Myllykoski, R. E. Looper, P. Koivunen, Cancer-associated 2-oxoglutarate analogues modify histone methylation by inhibiting histone lysine demethylases. J Mol Biol 430, 3081–3092 (2018).

18. J. Balss et al., Enzymatic assay for quantitative analysis of (D)-2-hydroxyglutarate. Acta Neuropathol 124, 883–891 (2012).

19. M. Batie, J. Druker, L. D’Ignazio, S. Rocha, KDM2 Family Members are Regulated by HIF-1 in Hypoxia. Cells 6, (2017).

20. J. Frost et al., Potent and selective chemical probe of hypoxic signalling downstream of HIF-alpha hydroxylation via VHL inhibition. Nat Commun 7, 13312 (2016).

21. A. Subramanian et al., Gene set enrichment analysis: a knowledge-based approach for interpreting genome-wide expression profiles. Proc Natl Acad Sci U S A 102, 15545–15550 (2005).

22. A. Liberzon et al., Molecular signatures database (MSigDB) 3.0. Bioinformatics 27, 1739–1740 (2011).

23. Y. Benita et al., An integrative genomics approach identifies Hypoxia Inducible Factor-1 (HIF-1)-target genes that form the core response to hypoxia. Nucleic Acids Res 37, 4587–4602 (2009).

24. A. Ortiz-Barahona, D. Villar, N. Pescador, J. Amigo, L. del Peso, Genome-wide identification of hypoxia-inducible factor binding sites and target genes by a probabilistic model integrating transcription-profiling data and in silico binding site prediction. Nucleic Acids Res 38, 2332–2345 (2010).

25. X. Zhou et al., Hypoxia induces trimethylated H3 lysine 4 by inhibition of JARID1A demethylase. Cancer research 70, 4214–4221 (2010).

26. G. Bellot et al., Hypoxia-induced autophagy is mediated through hypoxia-inducible factor induction of BNIP3 and BNIP3L via their BH3 domains. Molecular and cellular biology 29, 2570–2581 (2009).

27. V. E. Pedanou et al., The histone H3K9 demethylase KDM3A promotes anoikis by transcriptionally activating pro-apoptotic genes BNIP3 and BNIP3L. Elife 5, (2016).

28. G. D’Angelo, E. Duplan, N. Boyer, P. Vigne, C. Frelin, Hypoxia up-regulates prolyl hydroxylase activity: a feedback mechanism that limits HIF-1 responses during reoxygenation. J Biol Chem 278, 38183–38187 (2003).

29. M. Levin, M. Stark, Y. G. Assaraf, The JmjN domain as a dimerization interface and a targeted inhibitor of KDM4 demethylase activity. Oncotarget 9, 16861–16882 (2018).

30. D. B. Bekker-Jensen et al., An Optimized Shotgun Strategy for the Rapid Generation of Comprehensive Human Proteomes. Cell Syst 4, 587–599 e584 (2017).

31. Y. Huang, D. Chen, C. Liu, W. Shen, Y. Ruan, Evolution and conservation of JmjC domain proteins in the green lineage. Mol Genet Genomics 291, 33–49 (2016).

32. Y. Chang, J. Wu, X. J. Tong, J. Q. Zhou, J. Ding, Crystal structure of the catalytic core of Saccharomyces cerevesiae histone demethylase Rph1: insights into the substrate specificity and catalytic mechanism. Biochem J 433, 295–302 (2011).

33. B. Jia, K. Tang, B. H. Chun, C. O. Jeon, Large-scale examination of functional and sequence diversity of 2-oxoglutarate/Fe(II)-dependent oxygenases in Metazoa. Biochim Biophys Acta 1861, 2922–2933 (2017).

34. R. J. Klose et al., The retinoblastoma binding protein RBP2 is an H3K4 demethylase. Cell 128, 889–900 (2007).

35. C. Allan et al., OMERO: flexible, model-driven data management for experimental biology. Nature methods 9, 245–253 (2012).

36. K. Schumm, S. Rocha, J. Caamano, N. D. Perkins, Regulation of p53 tumour suppressor target gene expression by the p52 NF-kappaB subunit. The EMBO journal 25, 4820–4832 (2006).

37. Y. Liao, G. K. Smyth, W. Shi, The Subread aligner: fast, accurate and scalable read mapping by seed-and-vote. Nucleic Acids Res 41, e108 (2013).

38. F. Ramirez et al., deepTools2: a next generation web server for deep-sequencing data analysis. Nucleic Acids Res 44, W160–165 (2016).

39. H. Li et al., The Sequence Alignment/Map format and SAMtools. Bioinformatics 25, 2078–2079 (2009).

40. Y. Zhang et al., Model-based analysis of ChIP-Seq (MACS). Genome Biol 9, R137 (2008).

41. C. Zang et al., A clustering approach for identification of enriched domains from histone modification ChIP-Seq data. Bioinformatics 25, 1952–1958 (2009).

42. L. J. Zhu et al., ChIPpeakAnno: a Bioconductor package to annotate ChIP-seq and ChIP-chip data. BMC Bioinformatics 11, 237 (2010).

43. S. Durinck, P. T. Spellman, E. Birney, W. Huber, Mapping identifiers for the integration of genomic datasets with the R/Bioconductor package biomaRt. Nat Protoc 4, 1184–1191 (2009).

44. G. Yu, L. G. Wang, Q. Y. He, ChIPseeker: an R/Bioconductor package for ChIP peak annotation, comparison and visualization. Bioinformatics 31, 2382–2383 (2015).

45. A. Liberzon et al., The Molecular Signatures Database (MSigDB) hallmark gene set collection. Cell Syst 1, 417–425 (2015).

46. Y. Liao, G. K. Smyth, W. Shi, featureCounts: an efficient general purpose program for assigning sequence reads to genomic features. Bioinformatics 30, 923–930 (2014).

47. M. D. Robinson, D. J. McCarthy, G. K. Smyth, edgeR: a Bioconductor package for differential expression analysis of digital gene expression data. Bioinformatics 26, 139–140 (2010).

48. D. J. McCarthy, Y. Chen, G. K. Smyth, Differential expression analysis of multifactor RNA-Seq experiments with respect to biological variation. Nucleic Acids Res 40, 4288–4297 (2012).

49. R. Andersson et al., An atlas of active enhancers across human cell types and tissues. Nature 507, 455–461 (2014).

50. F. Consortium et al., A promoter-level mammalian expression atlas. Nature 507, 462–470 (2014).

51. M. Lizio et al., Update of the FANTOM web resource: high resolution transcriptome of diverse cell types in mammals. Nucleic Acids Res 45, D737–D743 (2017).

52. J. Liang et al., From a novel HTS hit to potent, selective, and orally bioavailable KDM5 inhibitors. Bioorg Med Chem Lett 27, 2974–2981 (2017).

53. C. Johansson et al., Structural analysis of human KDM5B guides histone demethylase inhibitor development. Nat Chem Biol 12, 539–545 (2016).

54. Z. Yang et al., Structure of the Arabidopsis JMJ14-H3K4me3 Complex Provides Insight into the Substrate Specificity of KDM5 Subfamily Histone Demethylases. Plant Cell 30, 167–177 (2018).

55. E. F. Pettersen et al., UCSF Chimera--a visualization system for exploratory research and analysis. J Comput Chem 25, 1605–1612 (2004).

56. F. Sievers et al., Fast, scalable generation of high-quality protein multiple sequence alignments using Clustal Omega. Mol Syst Biol 7, 539 (2011).

57. A. M. Waterhouse, J. B. Procter, D. M. Martin, M. Clamp, G. J. Barton, Jalview Version 2--a multiple sequence alignment editor and analysis workbench. Bioinformatics 25, 1189–1191 (2009).

