## Supplementary material for "Hypoxia induces rapid changes to histone methylation reprogramming chromatin for the cellular response"

Supplementary Materials for

### Hypoxia induces rapid changes to histone methylation marks and programs chromatin for the appropriate cellular response

Michael Batie, Julianty Frost, Mark Frost, James, W. Wilson, Pieta Schofield, and Sonia Rocha

**This PDF file includes:**

Materials and Methods
Supplementary Text
Figs. S1 to S13
References (34-57)

#### Materials and Methods

Cell lines and treatment

HeLa, HFF, 786-O, HEK293, HT1080, RCC4-HA and RCC4-HA-VHL cell lines were obtained from the American Type Culture Collection (ATCC). HeLa, HFF, HEK293, and HT1080 cell lines were maintained in Dulbecco's modified Eagle's medium (DMEM) (Sigma), supplemented with 2mM L- glutamine, 10% (v/v) Foetal Bovine Serum (FBS) (Gibco), 50units/mL penicillin (Lonza) and 50μg/mL streptomycin (Lonza). RCC4-HA and RCC4-HA-VHL cell lines were maintained as above with 400mg/mL G418 (Sigma). 786-O cells were maintained in Roswell Park Memorial Institute medium (RPMI) 1640 media (Gibco) supplemented with 2mM L-glutamine, 10% (v/v) FBS (Gibco), 50units/mL penicillin (Lonza) and 50μg/mL streptomycin (Lonza). Hypoxia treatments were performed in an InVivo 300 or InVivo2 300 hypoxia workstation (Ruskin) at 1% $O_2$, 5% $CO_2$, and 37°C unless stated otherwise. Dimethyloxalylglycine (DMOG) (Merck-Millipore) was dissolved in DMSO and used at a final concentration of 1mM. Desferroxamine mesylate (DFX) (Sigma) was dissolved in water and used at a final concentration of 0.2mM. VH298 (Sigma) was dissolved in DMSO and used at the final concentration of 0.1mM. Hydrogen peroxide ($H_2O_2$) (Sigma) was used at a final concentration of 0.5mM. Diethyl succinate (sigma) was used at a final concentration of 5mM. Diethyl fumarate (sigma) was used at a final concentration of 0.1mM. N-acetyl-cysteine (NAC) (Sigma) was dissolved in water and used at a final concentration of 0.5mM. Octyl-R-2HG (Sigma) was dissolved in DMSO and used at the final concentration of 5mM.

siRNA and DNA Transfection

siRNA transfections were performed using Interferin (Peqlab following manufacturer's instructions. For DNA transfections, in 6 well plates, 1μg of plasmid DNA, 0.12M CaCl2 and

HEPES buffered saline (0.156M NaCl, 0.375M Na2HPO4, 10mM HEPES) were mixed in water at a final volume of 400uL and added to cells. Cell culture media was changed 24H following transfection and cells were harvested 48H following transfection.

siRNAs

siRNA oligonucleotides were purchased from MWG and used in a final concentration of 27nM. The following primary siRNA oligonucleotides were used:

Control: 5'-CAGUCGCGUUUGCGACUGG-3' (*19*)

HIF-1β_1: 5'-GGUCAGCAGUCUUCCAUG-3' (*19*)

HIF-1β_2: 5'- GAAAGAAACAUGUGAGUAA-3' (*19*)

KDM2B: 5'-CCACUGCAAGUCUAGCACA-3' (*19*)

KDM5A#1: 5'-GAAGAAUUCUAGCCAUACA-3'

KDM5A#2: 5'-GGAAAUACCCAGAGAAUGA-3'

KDM5B: 5'- UAAGUUAGUUGCAGAAGAA-3'

KDM5C: 5'- GAAGAUUGUUGGUGUGUAA-3'

Plasmids

pcDNA3/HA-FLAG-RBP2(KDM5A) was a gift from William Kaelin (Addgene plasmid # 14800) (*34*). The following mutants were made on the KDM5A plasmid using site directed mutagenesis: T30A S34A, V109H, M297L, S464C, D1633N.

Immunoblots

Cells were lysed in Radio Immunoprecipitation Assay (RIPA) buffer (50 mM Tris-HCl (pH 8), 150mM NaCl, 1% (v/v) NP40, 0.25% (w/v) Na-deoxycholate, 0.1% (w/v) SDS, 10Mm NaF, 2mM

$Na_3VO_4$ and 1 tablet/10 mL Complete, Mini, EDTA-free protease inhibitors (Roche)) and centrifuged for 10min at 13000rpm, 4°C, and supernatants were collected. For acid extraction of histones, pellets from RIPA buffer lysed cells were resuspended in 400µL of buffer W (50mM Tris-HCl (pH 7.4) and 13mM EDTA) and centrifuged for 10min at 13000rpm, 4°C. The pellets were resuspended in 200µL of 0.4N $H_2SO_4$ and incubated on ice for 90min. The samples were centrifuged for 10min at 13000rpm, 4°C and the supernatant was mixed with 1mL acetone and incubated at -20°C overnight. The samples were centrifuged for 10min at 13000rpm, 4°C. The pellet was resuspended in 1mL Acetone and centrifuged for 10min at 13000rpm, 4°C. The pellets were then air dried for 10min then resuspended in 50µL of 4M urea and stored at -80°C. SDS-PAGE and immunoblots were carried out using standard protocols. Image J was used for quantification of immunoblots, with results expressed as mean and SEM band intensity normalised to β-Actin (and H3if appropriate) for a minimum of three independent experiments. The following primary antibodies were used immunoblotting: HIF 1α (610958, BD Biosciences), HIF 1β (3718, Cell Signaling), β-Actin (3700, CST)/ (60009-1-Ig, Proteintech), KDM5A (3876, Cell Signaling), H3K4me3 (9751, Cell Signaling), H3K9me3 (9754, Cell Signaling)/ (13969, Cell Signaling), H3K27me3 (9733, Cell Signaling), H3K36me2 (2901, Cell Signaling), H3K36me3 (4909, Cell Signaling), H3 (10809, Santa Cruz)/ (3638, Cell Signaling), BNIP3L (12396, Cell Signaling), LC3B (3868, Cell Signaling), p62 (610832,BD Biosciences). Phospho-AKT (Ser473) (4606, Cell signaling) BNIP3 (Ab10433, Abcam).

Immunofluorescence

Cells were plated onto sterilized glass coverslips (VWR 19mm) in 35mm plates at a density of $1.2x10^5$. 72H later cells were fixed to coverslips in 3.7% paraformaldehyde (w/v) in PBS (pH 6.8) for 15min at 37°C inside cell incubator or hypoxia workstation, washed with PBS and permeabilized in PBS-0.1% Triton X-100 (w/v) for 10mins at room temperature. Alternatively, cells were fixed in 100% methanol for 5min at -20°C. Cells were then washed with PBS and blocked with 1% donkey serum (v/v) in PBS 0.05% Tween (v/v) for 30min at room temperature. Cells were placed in a humidified chamber, incubated with primary antibody diluted in 1% donkey serum (v/v) in PBS 0.05% Tween (v/v) overnight, washed 3x 5min in PBS, incubated for 1H with secondary antibody in 1% donkey serum (v/v) in PBS 0.05% Tween (v/v) and washed 3x 5min in PBS. Nucleus staining was performed by incubation with 33.3μM Hoechst (Sigma) for 2min. Coverslips were then washed in water and mounted onto slides (VWR) using mounting medium (H-1000; Vector labs) and sealed with nail varnish. Cells were analysed and images were acquired using a DeltaVision microscope. Images were deconvolved and analysed using OMERO client software (Open Microscopy Environment (*35*)). The following primary antibodies were used for immunofluorescence: H3K4me3 (9751, Cell Signaling), H3K36me3 (4909, Cell Signaling).

Cellular Reactive Oxygen Species Detection

Cellular ROS were measured using CellRox staining (Thermo Fisher) following the exposure to 1 hour 1% $O_2$, according to the manufacturer's instructions. Cells were fixed and imaged using epifluorescence microscopy (Zeiss). Images were quantified using OMERO client software (*35*).

mRNA expression analysis by qPCR

Total RNA from cells was extracted using peqGOLD total RNA kit (Peqlab) and converted to cDNA using Quantitect Reverse Transcription Kit (Qiagen). Brilliant II Sybr green kit (Statagene/Agilent), and specific MX3005P 96 well semi-skirted plates, were used to analyse samples by qPCR on the MX3005P qPCR platform (Stratagene/Agilent). qPCR results are expressed normalised to β-Actin and relative to a calibrator condition, as mean and SEM of a minimum of three independent experiments. The following primers were used for qPCR:

β-Actin

F: 5’-CTGGGAGTGGGTGGAGGC-3’

R: 5’-TCAACTGGTCTCAAGTCAGTG-3’

KDM2B

KDM2B (Qiagen QT00087640)

KDM5A

F: 5’-GAGCTGTGTTCCTCTTCCTAAA-3’

R: 5’-CCTTCGAGACCGCATACAAA-3’

KDM5B

F: 5’-GCCCTCAGACACATCCTATTC-3’

R: 5’-AGTCCACCTCATCTCCTTCT-3’

KDM5C

F: 5’-ACAGAAGGAGAAGGAGGGTAT-3’

R: 5’-CACACACAGATAGAGGTTGTAGAG-3’

ENO1

F: 5’-CCTGGCATGGATCTTGAGAA -3’

R: 5’-TACGTTCACCTCGGTGTCTG-3’

CA9

F: 5’-CTTTGCCAGAGTTGACGAGG-3’

R: 5’-CAGCAACTGCTCATAGGCAC-3’

LOX

F: 5’-AGGCTGCCCTCTATCATT-3’

R: 5'-GTTTCACGGCTGCCTTAT-3'

KLF10

F: 5'-CTGTTGTGTTCATGGGCACA-3'

R: 5'-CTGTTGTGTTCATGGGCACA-3'

STAG2

F: 5'-GGTCCAAATGCCAACCTTGT-3'

R: 5'-CTCAGTAGCACAGTCCCACA-3'

BNIP3L

F: 5'-GTGGAAATGCACACCAGCAG-3'

R: 5'-CTTGGGTGGAATGTTTTCGG-3'

BAP1

F: 5'- GGAGGTAGAGAAGAGGAAGAA-3'

R: 5'- GAGCCAGCATGGAGATAAAG -3'

<u>ChIP qPCR</u>

Chromatin Immunoprecipitation (ChIP) was performed using an adaptation from (*36*). Protein was cross-linked to chromatin by addition of formaldehyde to a final concentration of 1% for 10min at 37°C inside cell incubator or hypoxia workstation. Glycine was added to a final concentration of 0.125M for 5min at room temperature to quench excess formaldehyde. Cells were washed 2x in DPBS, harvested in 400μL of ChIP lysis buffer (50mM Tris-HCl (pH 8.1), 1% (w/v) SDS, 10mM EDTA and 1 Complete, Mini, EDTA-free protease inhibitors (Roche)) per 150mm plate and incubated on ice for 10min. Samples were then sonicated for 8 cycles at 4°C for 15sec with a 30sec gap between each sonication at 50% amplitude (Sonics Vibra-Cell # VCX130). Samples were centrifuged for 10min at 13000rpm, 4°C and supernatants were collected. 10μL of samples were used for input and stored at -20°C. 100µL of samples were diluted 10-fold in dilution buffer (20mM Tris-Hcl (pH 8.1), 1% (v/v) Triton X-100, 2mM EDTA and 150mM NaCl). Diluted samples were pre-cleared with 2µg of sheared salmon sperm DNA and 20µL of protein G-Sepharose (50% slurry) (Generon) on a rotating wheel at 4°C for 2H. Immunoprecipitations were performed by incubating 1mL of samples with antibodies and 0.1% (v/v) Brij 35 overnight on a rotating wheel at 4°C. Immune complexes were captured by incubation with 2µg of sheared salmon sperm DNA and 30µL of protein G-Sepharose (50% slurry) on a rotating wheel at 4°C for 1H. Immunoprecipitates were washed sequentially on a rotating wheel at 4°C for 5min each with Wash Buffer 1 (20mM Tris-HCl (pH 8.1), 0.1% (w/v) SDS, 1% (v/v) Triton X-100, 2mM EDTA and 150mM NaCl), Wash Buffer 2 (20mM Tris-HCl (pH 8.1), 0.1% (w/v) SDS, 1% (v/v) Triton X-100, 2mM EDTA and 500mM NaCl), and Wash Buffer 3 (10mM Tris-HCl (pH 8.1), 0.25M LiCl, 1% (v/v) NP-40, 1% (w/v) Na-deoxycholate and 1mM EDTA), 2x with TE buffer (10mM Tris-HCl (pH 8.0) and 1mM EDTA ) and eluted with 120µL of Elution Buffer (1% (w/v) SDS and 0.1M Na-bicarbonate). Crosslinks were reversed by incubation of samples (including inputs) with 0.2M NaCl overnight on a thermocycler at 300rmp, 65°C. Proteins were digested by incubation of samples with 40mM Tris-HCl (pH6.5), 10mM EDTA and 20µg Proteinase K for 1H at 45°C. DNA was purified using a PCR-product purification kit according to manufacturer's instructions (NBS Biologicals #NBS363). 3µL of DNA was used for qPCR analysis. ChIP qPCR results are expressed relatively to input material and a calibrator condition, as mean and SEM of a minimum of three independent experiments. The following antibodies were used for ChIP qPCR: H3K4me3 (39915, Active Motif), H3K9me3 (39161, Active Motif), H3K36me3 (61101, Active Motif), H3 (61475, Active Motif), Rabbit IgG (I5006, Sigma), Mouse IgG (I5381, Sigma). The following primers were used for ChIP qPCR:

ENO1

F: 5’-CTCTCTAGTCCATGTCATGTTG-3’

R: 5’-GTTCACAAGAATCCAGGTAGG-3’

LOX

F: 5’-CTTTCATCTCCACAGTGTCTAC-3’

R: 5’-GCCTTCTGAGTGCTTTCTC-3’

KLF10

F: 5’-TGCCTTCGTGTTGAAATCC-3’

R: 5’-GGGCGCGATTATGCAATTA-3’

STAG2

F: 5’-AAAAATGTCTCAGATTTTACTTCTTGT-3’

R: 5’-CTCTTGTTAGTTTCTCTCCCATCC-3’

BNIP3L

F: 5’-GTGCCTTGCTTCTCCTTT-3’

R: 5’-GGCTGCTCTCAGCTTTAC-3’

CA9

F: 5’-CTTCTGGTGCCTGTCCATC-3’

R: 5’-ATCCTCCTCTCTGGGTGAAT-3’

β-Actin

F: 5’- GCGGTGCTAGGAACTCAAA-3’

R: 5’- TACTCAGTGGACAGACCCAA-3’

KDM2B

F: 5’- CCTAGTAAAGGAGTCCACAG-3’

R: 5’- CCATACCTATAAGGACTGCC-3’

BAP1

F: 5’- TCTGAAGGTCGTAGATCTCCTC-3’

R: 5’- CTGTCCTTCCCTACTGCTTTC-3’

ChIP-Sequencing

The ChIP protocol described above was followed however immunoprecipitation and input volumes were upscaled 6x, sheared salmon sperm DNA and protein G-Sepharose (50% slurry) volumes were upscaled x2 and following final wash of immunoprecipitates with TE buffer, immune complex elution, reverse crosslinks, protein digestion and DNA elution was performed using a magnetic DNA purification kit according to manufacturer’s instructions (Diagenode C03010011). 10ng of DNA from each ChIP sample and corresponding input (three replicates per condition) were used in single end DNA library construction and sequenced on a HiSeq 2000 platform (Illumina) at the University of Dundee Genome Sequencing Unit.

ChIP-Sequencing data analysis

Fastq data files were passed through the quality control tool FASTQC and aligned to the human genome annotation hg19_73 using the subread package (*37*). The data were then converted to Bam and then RPKM normalised bigwig file formats using github packages (*38, 39*) H3K4me3 peaks for individual replicates and merged bam files for each condition (normoxia (21% $O_2$) and hypoxia (1H, 1% $O_2$)) were called using the MACS2 peak caller (*40*) in narrow peak mode with the IP sample bam files set as treatment and corresponding input bam files set as control. Peaks called in 3 out of 3 replicates per condition were designated as high stringency peaks. To call H3K4me3 peaks enriched in hypoxia over normoxia , the merged hypoxia bam file was used as the treatment and merged normoxia bam file as the control in MACS2 peak caller narrow peak mode. To call H3K4me3 peaks enriched in normoxia over hypoxia, the merged normoxia bam file was used as the treatment and the merged hypoxia bam file as the control in MACS2 peak caller narrow peak mode. An FDR of 0.05 was used as the cut off for MACS2 peak calls. H3K36me3 peaks for individual replicates and merge bam files for each condition were called using the SICER peak caller (*41*) with the IP sample bam files set as treatment and corresponding input bams file set as control. SICER is a peak caller which specializes in calling broad peaks and was deemed more suitable than MACS2 for calling H3K36me3 peaks. To call H3K36me3 peaks enriched in hypoxia over normoxia, the merged hypoxia bam file was used as the treatment and merged normoxia bam file as the control. To call H3K36me3 peaks enriched in normoxia over hypoxia, the merged normoxia bam file was used as the treatment and the merged hypoxia bam file as the control. FDR of 0.01 was used as the cut off for all SICER calls with a fold change cut off of 0.2 for peaks called as enriched in one condition over another. H3K4me3 and H3K36me3 1H, 1% $O_2$ upregulated downregulated peaks were further filtered to only include high stringency peaks (peaks called in 3 out of 3 replicates) for the condition they are the enriched in. R Bioconductor packages were used for overlapping peaks (*42*), and identification of the nearest gene for each peak and genomic annotation assignment (*43, 44*). A Github package (*38*) was used to generate heat maps of RPKM and RPM normalised read counts to transcription start sites of protein coding genes. Coverage tracks for RPKM normalised reads were generated using the IGV genome browser. Pathway enrichment analysis was performed using GSEA MSigDB online tool with hallmark genes (*21, 22, 45*) with a 0.05 FDR and 0.05 p value cut off. The ChIP sequencing data were deposited to the NCBI GEO (GSE120339).

RNA-sequencing and data analysis

HeLa cells were seeded in 35 mm dishes one day prior to treatments with hypoxia (1% $O_2$), for 16 hours. RNA was extracted as described above and DNAse treatment was included according to manufacturer's instructions. Experiments were performed in six replicates. RNA samples used in library preparation and running of the samples on the NextSeq 500 platform. RNA ERCC ExFold RNA Spike-In Mix (Mix1 and Mix2) was distributed throughout the RNA-Seq experiment.

The total RNA integrity was determined using an Agilent 2200 TapeStation (Agilent Technologies, Edinburgh) on a standard sensitivity RNA ScreenTape according to instrument guidelines. The concentration of each sample was measured fluorometrically, using the QuBit 3.0 system on a standard sensitivity protocol (Thermo Fisher, UK). All samples were diluted to 200 ng/µl for a total input of 1 µg into the RNA-seq protocol. RNA ERCC ExFold RNA Spike-In Mix (Mix1 and Mix2) was distributed throughout the RNA-Seq experiment according to manufacturer's protocol (Ambion, 4456739 and 4456740, Rev D).

The TruSeq Stranded Total RNA with Ribo-ZeroTM Human/Mouse/Rat kit from Illumina (Illumina, UK) was utilised according to manufacturer's instruction, followed by verification of each sample library through Qubit 3.0 and Agilent 2200 TapeStation. Libraries were pooled to 10 nM, allowing 15 samples per run on the NextSeq 500 platform. The subsequent pooled library was verified on TapeStation and QuBit before dilution to 4 nM for inclusion into the sequencing protocol. To achieve the highest performance on the Illumina platform, the library pools were loaded at 1.8 pM with a 1% PhiX spike in for optimum cluster deposition. Paired-end Illumina sequencing was performed on the NextSeq 500 system using the High-Output v2, 150-cycle kit (76x76 bp PE sequencing). FastQ files were generated using the bcl2fastq conversion software before further analysis.

FastQ files containing reads were quality-controlled with *fast*QC (http://www.bioinformatics.bbsrc.ac.uk/projects/fastqc). The raw sequence reads from each replicate were aligned to the Ensembl human genome GRCh37 and ERCC sequence with STAR. The aligned reads were combined and number of read for each gene was counted with subreadfeatureCounts pipeline (*37, 46*). Contaminants were removed manually. Differential gene expression analysis was then performed by the R-package *edgeR* according to its user guide (*47, 48*).

Data mining/publicly available databases

The HIF target gene list was adapted from (*22*). Hypoxia induced and repressed genes across several cell types (Conserved hypoxia induced and repressed) gene lists provided by Dr Luis del Peso Ovalle (The Autonomous University of Madrid, Spain) (*23*). HeLa hypoxia induced and repressed gene lists were downloaded from NCBI GEO. (GSE120675). The following sequencing datasets from the ENCODE project (40, 41) were downloaded from NCBI GEO; HeLa S3 H3K4me3 (GSM945201), HeLa S3 H3K36me3 (GSM733711), HeLa S3 H3K27Ac (GSM733684) and HeLa S3 DNaseI hypersensitivity (GSM816633). Human Enhancer-transcription start site associations from FANTOM5/ Human Transcribed Enhancer Atlas were downloaded from the FANTOM5 web resource (*49-51*).

Colony formation assay

24H after transfection, cells were counted and plated on to 6 well plates at the density of 5000 cells per well. Cells were allowed to grow for 7 days prior to staining with Crystal Violet. For hypoxia treatment, plates were places at 1% $O_2$ 4 days after platting. Plates were imaged and colonies counted using Image J. Results are expressed as mean and SEM colony numbers of a minimum of three independent experiments.

Proliferation assay

24H after transfection, cells were counted and plated on to 6 well plates at the density of 40000 cells per well. Cells were counted 24 hours and for hypoxia treatment placed at 1% $O_2$. Cells were 48 hours later using a haemocytometer in triplicate. Results are expressed as mean and SEM of total cell counts in each day as, of a minimum of three independent experiments.

SubG1 analysis

Cells were treated and processed as in described for the Proliferation assays. After cell counting, cells were fixed in 70% ethanol and frozen at -20C for at least 1 day prior to staining and flow cytometry analysis. This was conducted using Cell Cycle analysis Kit from Guava (Millipore 4500-0220) and analysed in a Guava® easycyte HT (Millipore) cytometer.

Protein structural alignment/analysis

The structures were downloaded from the Protein Data Bank for KDM5A (PDB: 5v9p) (*52*), KDM5B (PDB: 5a3p) (*53*), KDM5C (PDB: 5fwj) (*53*), and the JMJ14 catalytic domain in complex with NOG and H3K4me3 peptide (PDB: 5yko) (*54*). UCSF Chimera (*55*) was used for the structure alignment and image rendering.

Sequence alignment

The KDM5A, KDM5B, and KDM5C amino acid sequences were aligned using Clustal Omega (*56*) and Jalview (*57*) was used for figure creation.

2-HG measurements

Intracellular 2-HG was measured using the 2-HG fluorimetric assay (Sigma) following the manufacturer's instructions, using HeLa, HFF and HT1080 cells.

KDM copy number analysis

Quantitative proteomic analysis of KDM family members was performed by analyzing copy number in HeLa cells in datasets from publicly available work (*29*) deposited on PRIDE (https://www.ebi.ac.uk/pride/archive/).

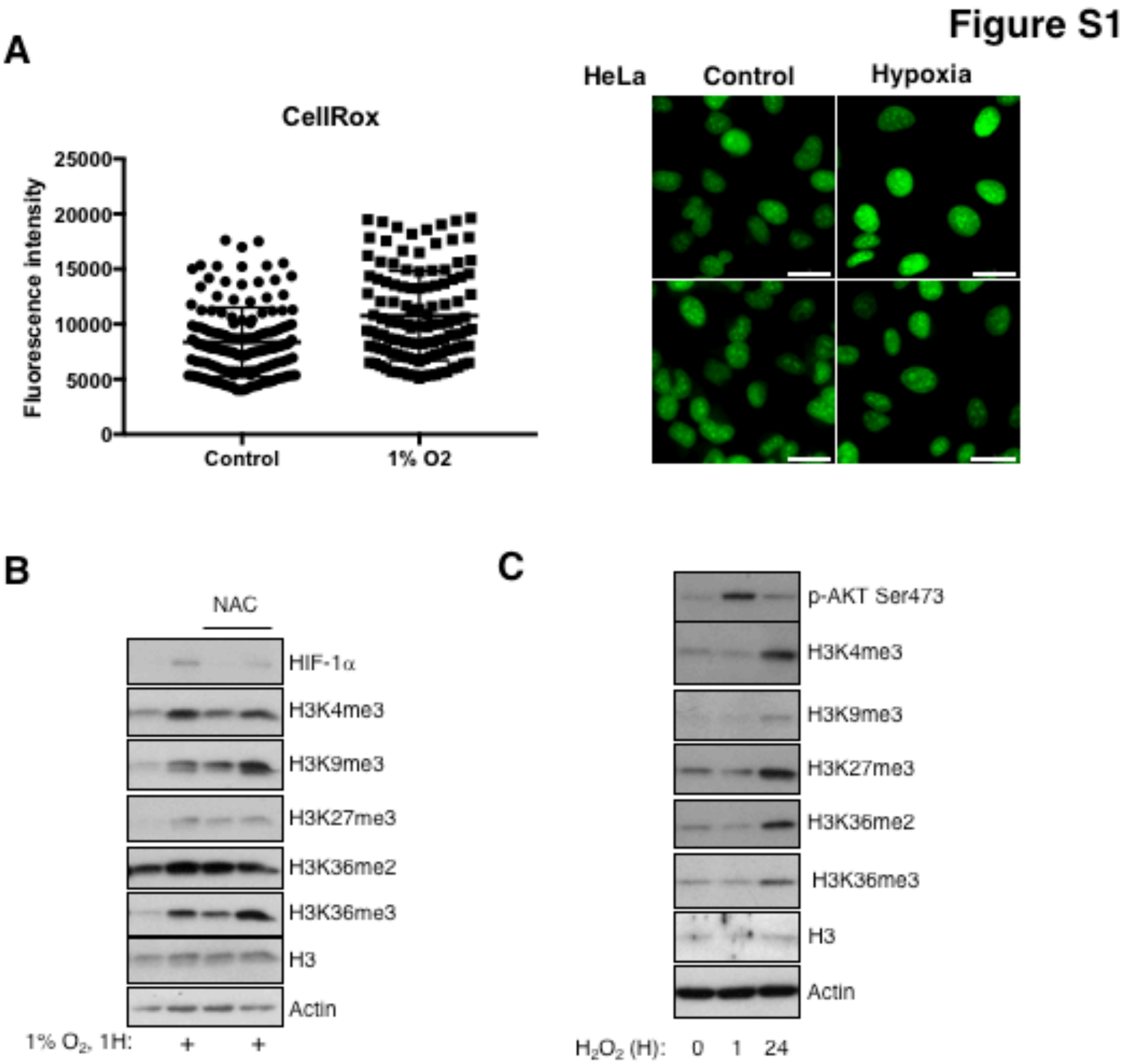
Figure S1
A
CellRox
Fluorescence intensity
25000
20000
15000
10000
5000
0
Control
1% O2
HeLa
Control
Hypoxia
B
NAC
HIF-1α
H3K4me3
H3K9me3
H3K27me3
H3K36me2
H3K36me3
H3
Actin
1% O2, 1H:
+
+
C
p-AKT Ser473
H3K4me3
H3K9me3
H3K27me3
H3K36me2
H3K36me3
H3
Actin
H2O2 (H):
0
1
24

**Fig. S1.**

**ROS are not involved in early hypoxia induced histone methylation.** CellRox quantification for HeLa cells exposed to 1% $O_2$ for 1 hour. Representative images are shown. Scale bar 20μM (**A**). Immunoblot analysis for histone methylation marks in HeLa cells following treatment with 1% $O_2$ for 1 hour with or without pre-treatment with NAC for 1 hour (**B**). Immunoblot analysis for histone methylation marks in HeLa cells following treatment with 500μM $H_2O_2$ for the indicated periods of time (**C**).

### Figure S2

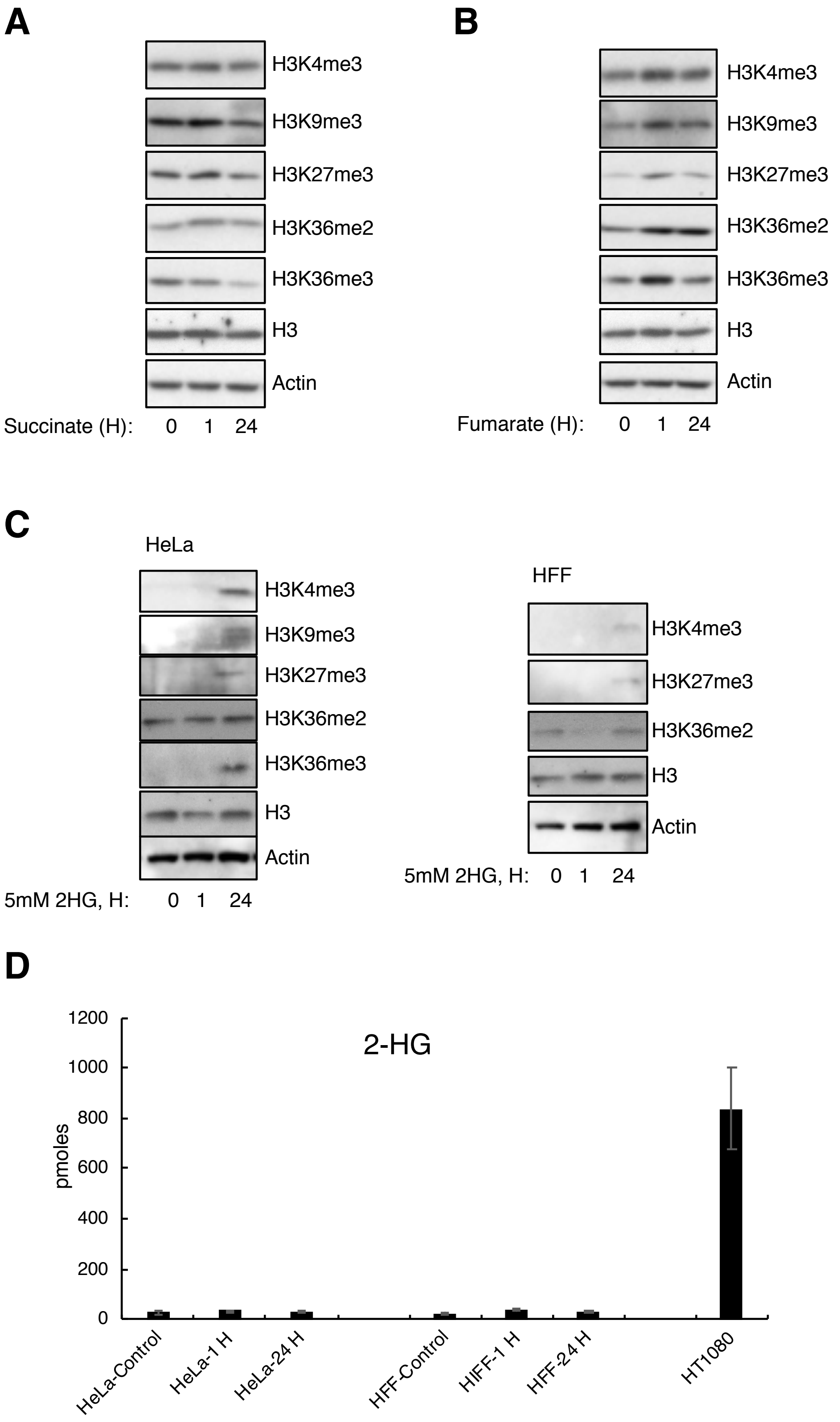
A
H3K4me3
H3K9me3
H3K27me3
H3K36me2
H3K36me3
H3
Actin
Succinate (H): 0 1 24
B
H3K4me3
H3K9me3
H3K27me3
H3K36me2
H3K36me3
H3
Actin
Fumarate (H): 0 1 24
C
HeLa
H3K4me3
H3K9me3
H3K27me3
H3K36me2
H3K36me3
H3
Actin
5mM 2HG, H: 0 1 24
HFF
H3K4me3
H3K27me3
H3K36me2
H3
Actin
5mM 2HG, H: 0 1 24
D
2-HG
pmoles
0
200
400
600
800
1000
1200
HeLa-Control
HeLa-1 H
HeLa-24 H
HFF-Control
HFF-1 H
HFF-24 H
HT1080

**Fig. S2.**

**Metabolite involvement in histone methylation.** Immunoblot analysis for histone methylation marks in HeLa cells following treatment with succinate (**A**), fumarate (**B**), or 2-HG (HFF cells were also analyzed) (**C**) Measurement of intracellular 2-HG in HeLa, HFF and HT1080 cells (**D**).

### Figure S3

**A**

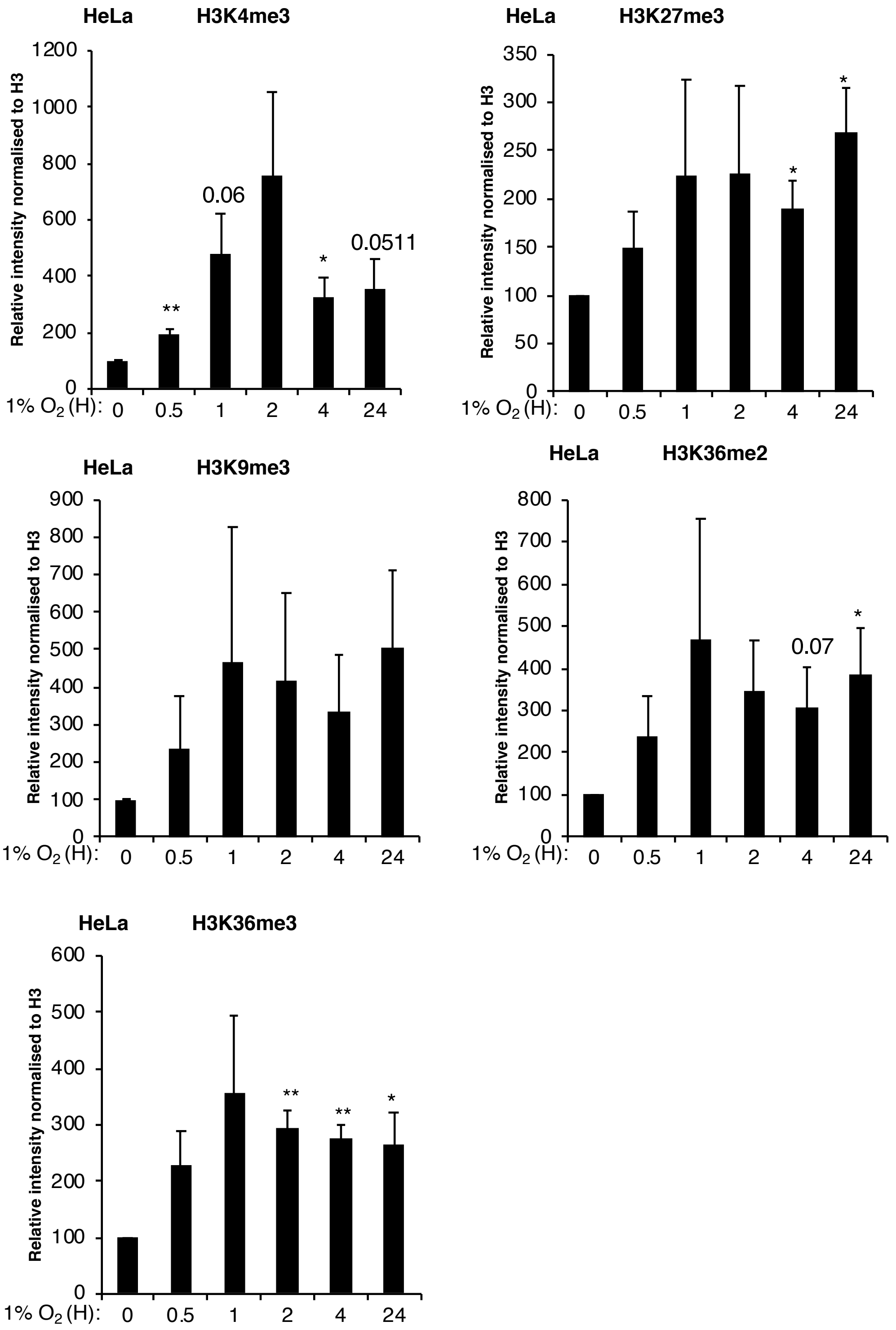
HeLa
H3K4me3
Relative intensity normalised to H3
1200
1000
800
600
400
200
0
0.06
0.0511
**
*
1% O2 (H):
0
0.5
1
2
4
24
HeLa
H3K27me3
Relative intensity normalised to H3
350
300
250
200
150
100
50
0
*
*
1% O2 (H):
0
0.5
1
2
4
24
HeLa
H3K9me3
Relative intensity normalised to H3
900
800
700
600
500
400
300
200
100
0
1% O2 (H):
0
0.5
1
2
4
24
HeLa
H3K36me2
Relative intensity normalised to H3
800
700
600
500
400
300
200
100
0
0.07
*
1% O2 (H):
0
0.5
1
2
4
24
HeLa
H3K36me3
Relative intensity normalised to H3
600
500
400
300
200
100
0
**
**
*
1% O2 (H):
0
0.5
1
2
4
24

**Fig. S3.**

**Hypoxia increases chromatin bound histone methylation.**

Signal intensity from immunoblot analysis of histone methylation on purified histone extracts from Hela cells exposed to 1% $O_2$ for the indicated time points (**A**). Graph depicts mean and SEM from a minimum of three independent experiments. Student's t test determined statistical significance. *$p < 0.05$, **$p < 0.01$, ***$p < 0.001$.

#### Figure S4

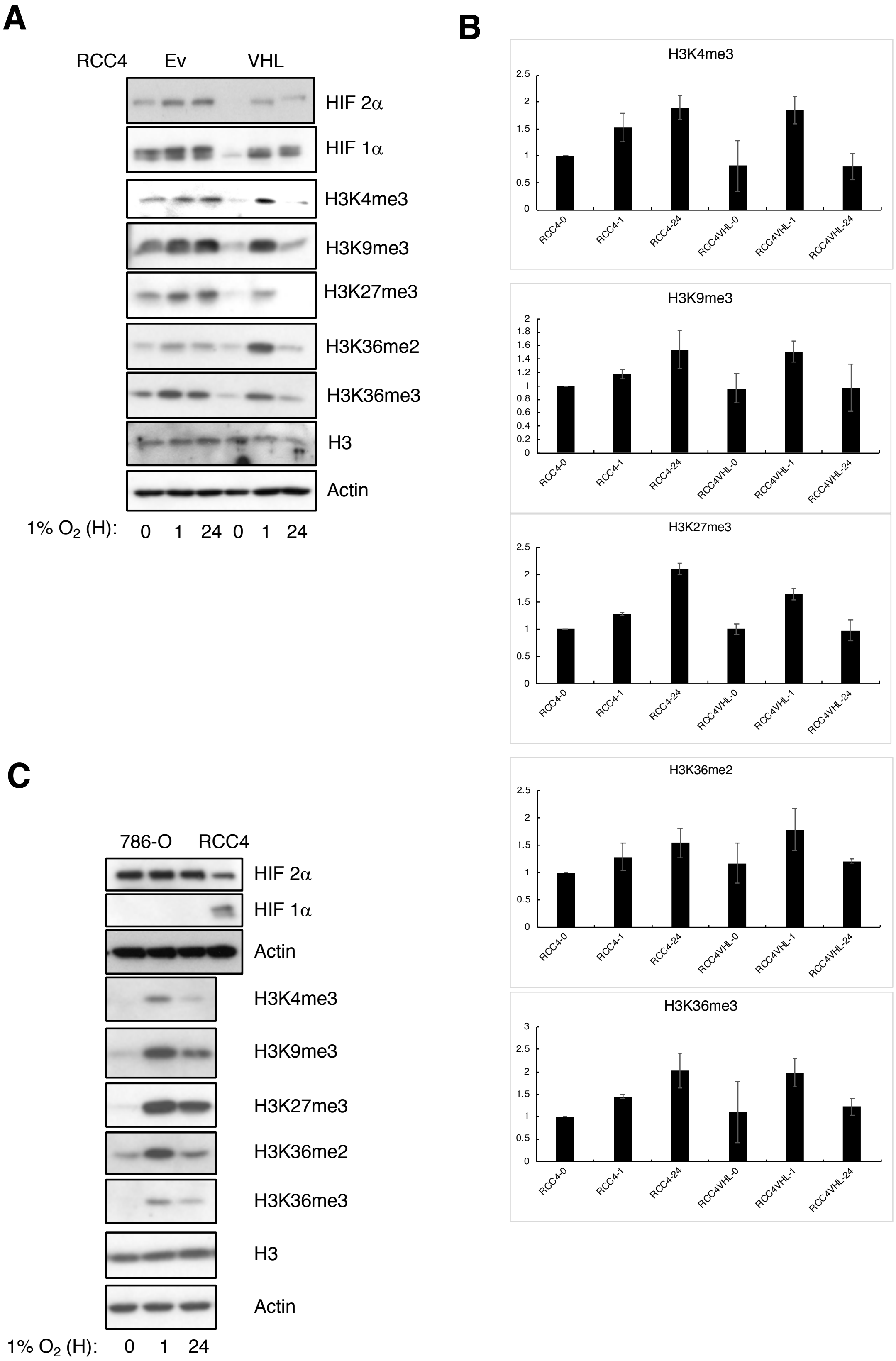
A
RCC4 Ev VHL
HIF 2α
HIF 1α
H3K4me3
H3K9me3
H3K27me3
H3K36me2
H3K36me3
H3
Actin
1% O2 (H): 0 1 24 0 1 24
B
H3K4me3
H3K9me3
H3K27me3
H3K36me2
H3K36me3
RCC4-0
RCC4-1
RCC4-24
RCC4VHL-0
RCC4VHL-1
RCC4VHL-24
C
786-O RCC4
HIF 2α
HIF 1α
Actin
H3K4me3
H3K9me3
H3K27me3
H3K36me2
H3K36me3
H3
Actin
1% O2 (H): 0 1 24

**Fig. S4.**

**Hypoxia increases histone methylation in cells lines expressing and not expressing HIF 1α.** Immunoblot analysis of histone methylation and HIF 1α in RCC4 cells with and without reconstituted VHL and exposed to 1% $O_2$ for the indicated time points (**A**). Quantification of histone marks using Image J (**B**). Immunoblot analysis of histone methylation in 786-O cells exposed to 1% $O_2$ for the indicated time points (**C**).

### Figure S5

**A**

| H3K4me3 peaks | 21% $O_2$ | 1H 1% $O_2$ |
|---|---|---|
| Rep1 | 34510 | 20974 |
| Rep2 | 21255 | 25433 |
| Rep3 | 39120 | 38992 |
| 3/3 | 14322 | 14062 |
| Merged | 42815 | 43249 |

**B**

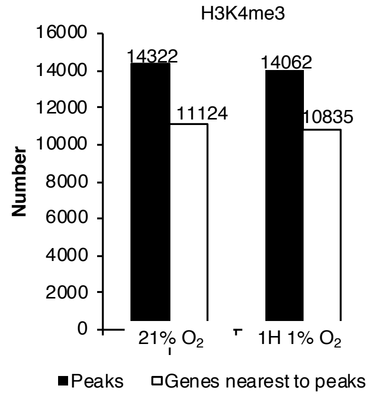


**C**

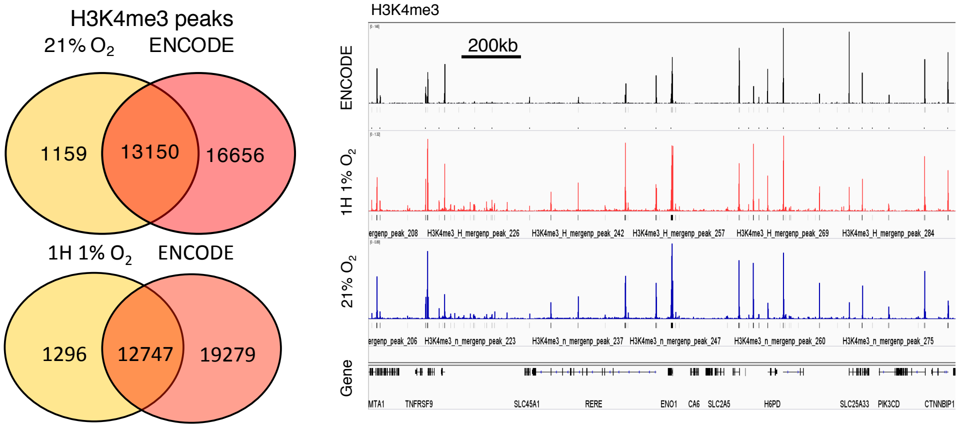


**D**

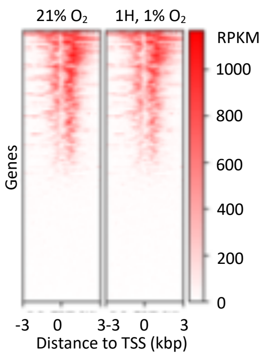


**E**

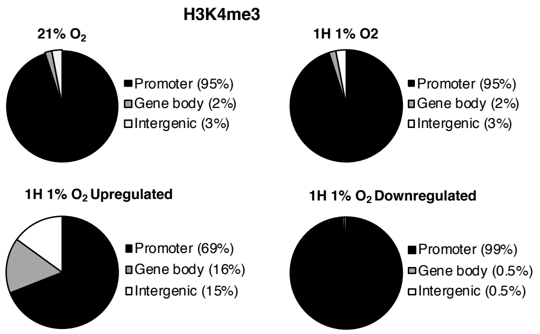


**F**

**Overlap of H3K4 1H 1% $O_2$ upregulated intergenic peaks with enhancers**

| Peaks | DNase HS sites (172580) | H3K27ac peaks (46470) | Both (32100) | None |
|---|---|---|---|---|
| 25 | 20 (80%) | 15 (60%) | 15 (60%) | 5 (20%) |

| Peaks | Fantom5 enhancers (32692) |
|---|---|
| 25 | 5 (25%) |

| Peak coordinates | Fantom5 enhancer coordinates | Fantom5 linked gene promoters |
|---|---|---|
| chr3:105,723,110-105,723,332 | chr3:105,722,948-105,723,182 | CBLB |
| chr5:148,188,828-148,189,046 | chr5:148,188,522-148,188,838 | FBXO38, ADRB2 |
| chr6:74,295,606-74,295,807 | chr6:74,295,249-74,295,764 | CGAS, EEF1A1, CD109 |
| chr9:21,677,715-21,677,893 | chr9:21,677,834-21,678,156 | ZNF217 |
| chr20:52,445,656-52,445,955 | chr20:52,445,816-52,446,339 | NA |

**Fig. S5.**

**H3K4me3 ChIP sequencing results.**

ChIP-Seq analysis of H3K4me3 in HeLa cells at 21% O2 (normoxia) or exposed to 1% $O_2$ oxygen for 1 hour (hypoxia). Number of ChIP-Seq peaks. (**A**). Number of high stringency (3/3 replicates) ChIP-seq peaks and genes nearest to peaks (**B**). Overlap of ChIP-Seq peaks with an ENCODE dataset for H3K4me3 and coverage tracks of these datasets (**C**). Heatmap of ChIP-Seq RPKM normalised read counts at transcription start sites of protein coding genes (**D**). Genomic annotation of ChIP-seq peaks, and differentially regulated peaks compared to normoxia (**E**). Analysis of intragenic H3K4me3 hypoxia upregulated peaks with enhancer markers (**F**).

**A** **Figure S6**

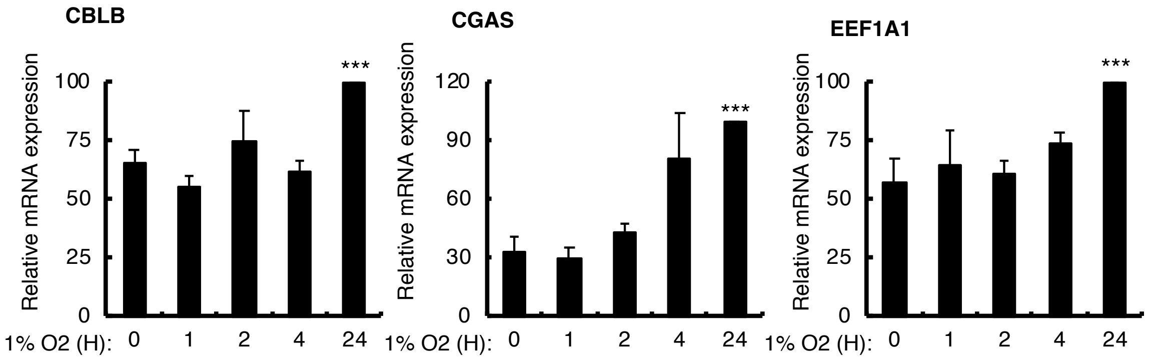
CBLB
Relative mRNA expression
100
75
50
25
0
***
1% O2 (H): 0 1 2 4 24
CGAS
Relative mRNA expression
120
90
60
30
0
***
1% O2 (H): 0 1 2 4 24
EEF1A1
Relative mRNA expression
100
75
50
25
0
***
1% O2 (H): 0 1 2 4 24

**Fig. S6.**

**Several genes associated with predicted enhancer identified with increased H3K4me3 peaks are hypoxia inducible.**

qPCR analysis of mRNA expression levels for the indicated genes in HeLa cells exposed to 1% $O_2$ for the indicated time points (**A**). Graph depicts mean and SEM from a minimum of three independent experiments. Student's t test determined statistical significance. *$p < 0.05$, **$p < 0.01$, ***$p < 0.001$.

### Figure S7

**A**

| H3K36me3 peaks | 21% $O_2$ | 1H 1% $O_2$ |
|---|---|---|
| Rep1 | 5102 | 5874 |
| Rep2 | 4707 | 3450 |
| Rep3 | 7574 | 6896 |
| 3/3 | 3829 | 3125 |
| Merged | 13864 | 12592 |

**B**

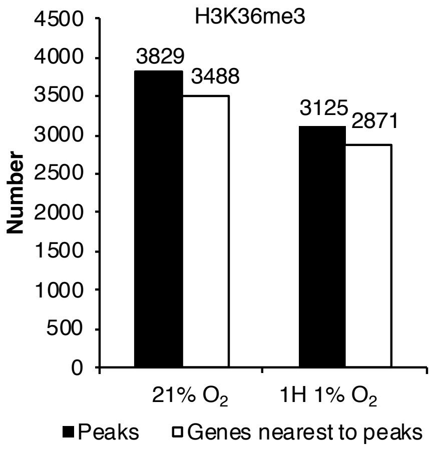


**C**

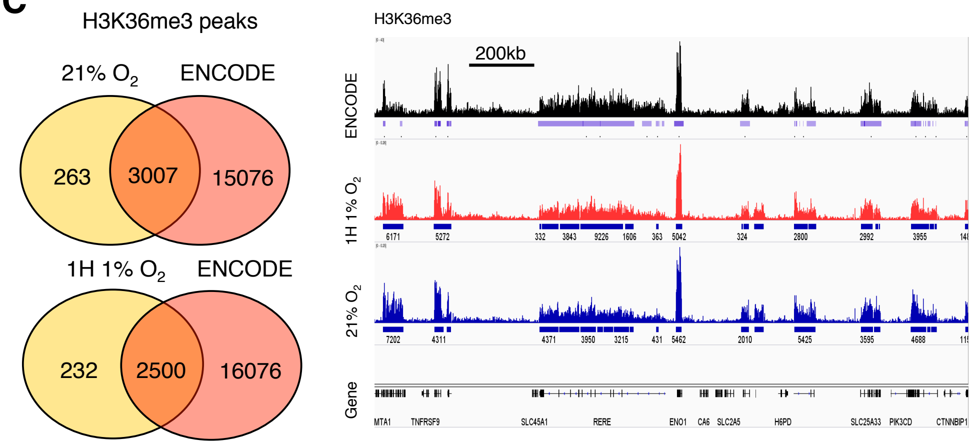


**D**

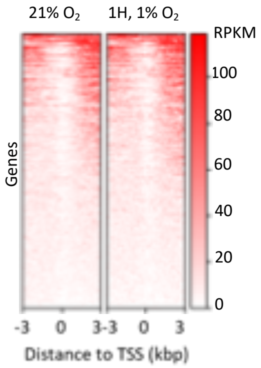


**E**

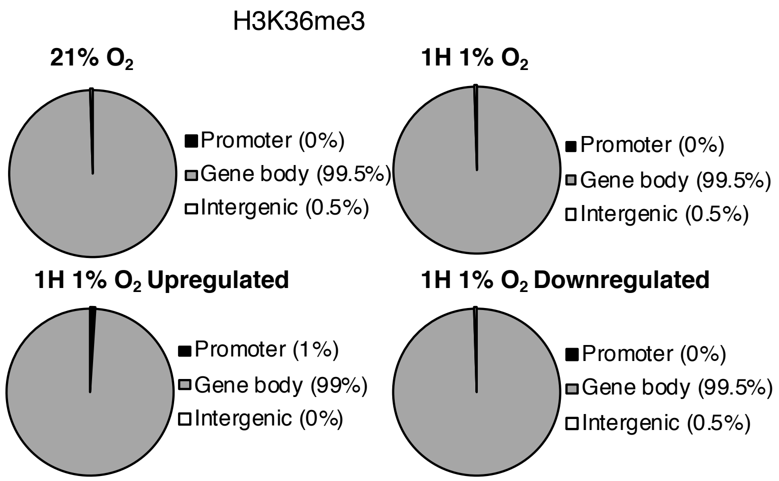

**Fig. S7.**
**H3K36me3 ChIP sequencing results.**
ChIP-Seq analysis of H3K4me3 in HeLa cells at 21% O2 (normoxia) or exposed to 1% $O_2$ for 1 hour (hypoxia). Number of ChIP-Seq peaks (**A**). Number of high stringency (3/3 replicates) ChIP-Seq peaks and genes nearest to peaks (**B**). Overlap of ChIP-seq peaks with an ENCODE dataset for H3K36me3 and coverage tracks of these datasets (**C**). Heatmap of ChIP-Seq RPKM normalised read counts at transcription start sites of protein coding genes (**D**). Genomic annotation of ChIP-seq peaks, and differentially regulated peaks compared to normoxia (**E**).

### Figure S8

## A

##### H3K4me3 1H 1% $O_2$ Downregulated

| Gene set | p value | Overlap |
|---|---|---|
| Mitotic spindle (200) | 1.73x10-6 | 11 |
| Oxidative phosphorylation (200) | 1.18x1-5 | 10 |
| Estrogen response late (200) | 7.3x10-5 | 9 |
| G2M (200) | 7.3x10-5 | 9 |
| DNA repair (150) | 5.61x10-5 | 8 |
| Estrogen response early (200) | 4.04x10-4 | 8 |
| Heme metabolism (200) | 4.04x10-4 | 8 |
| Myogeneis (200) | 4.04x10-4 | 8 |
| Interferon gamma response (200) | 8.52x10-3 | 6 |
| p53 pathway (200) | 8.52x10-3 | 6 |
| MYC targets (58) | 1.67x10-3 | 4 |
| Interferon α response (97) | 1.05x10-2 | 4 |
| Peroxisome (104) | 1.33x10-2 | 4 |
| PI3K/AKT mTOR signaling (105) | 1.37x10-2 | 4 |

-log10 (pvalue)

0-1.9 2-3.9 4-5.9 >6

## B

##### H3K36me3 1H 1% $O_2$ Downregulated

| Gene set | p value | Overlap |
|---|---|---|
| Response to UV radiation down (158) | 3.9x10-7 | 11 |
| MYC targets (200) | 2.46x10-5 | 10 |
| Oxidative phosphorylation (200) | 2.46x10-5 | 10 |
| Adipogenesis (200) | 1.39x10-4 | 9 |
| p53 pathway (200) | 1.39x10-4 | 9 |
| Mitotic spindle (200) | 7.08x10-4 | 8 |
| DNA repair (150) | 6.14x10-4 | 7 |
| Estrogen response late (200) | 3.19x10-3 | 7 |
| Bile acid metabolism (112) | 7.32x10-4 | 6 |
| Unfolded protein response (113) | 7.67x10-4 | 6 |
| Apical junction (200) | 1.26x10-2 | 6 |
| E2F targets (200) | 1.26x10-2 | 6 |
| Estrogen response early (200) | 1.26x10-2 | 6 |
| G2M (200) | 1.26x10-2 | 6 |
| Interferon gamma response (200) | 1.26x10-2 | 6 |
| Xenobiotic metabolism (200) | 1.26x10-2 | 6 |
| TGF β signaling (64) | 1.75x10-3 | 4 |
| Interferon α response (97) | 1.4x10-2 | 4 |

-log10 (pvalue)

0-1.9 2-3.9 4-5.9 >6

**Fig. S8.**

**Gene group association analysis for hypoxia downregulated H3K4me3 and H3K36me3 ChIP-seq peak genes.**

Gene group association analysis showing significant enrichment of gene set signatures (MsigDB) for H3K4me3 (**A**) and H3K36me3 (**B**) hypoxia downregulated ChIP-Seq peak genes. Hypergeometric distribution determined statistical significance.

### Figure S9

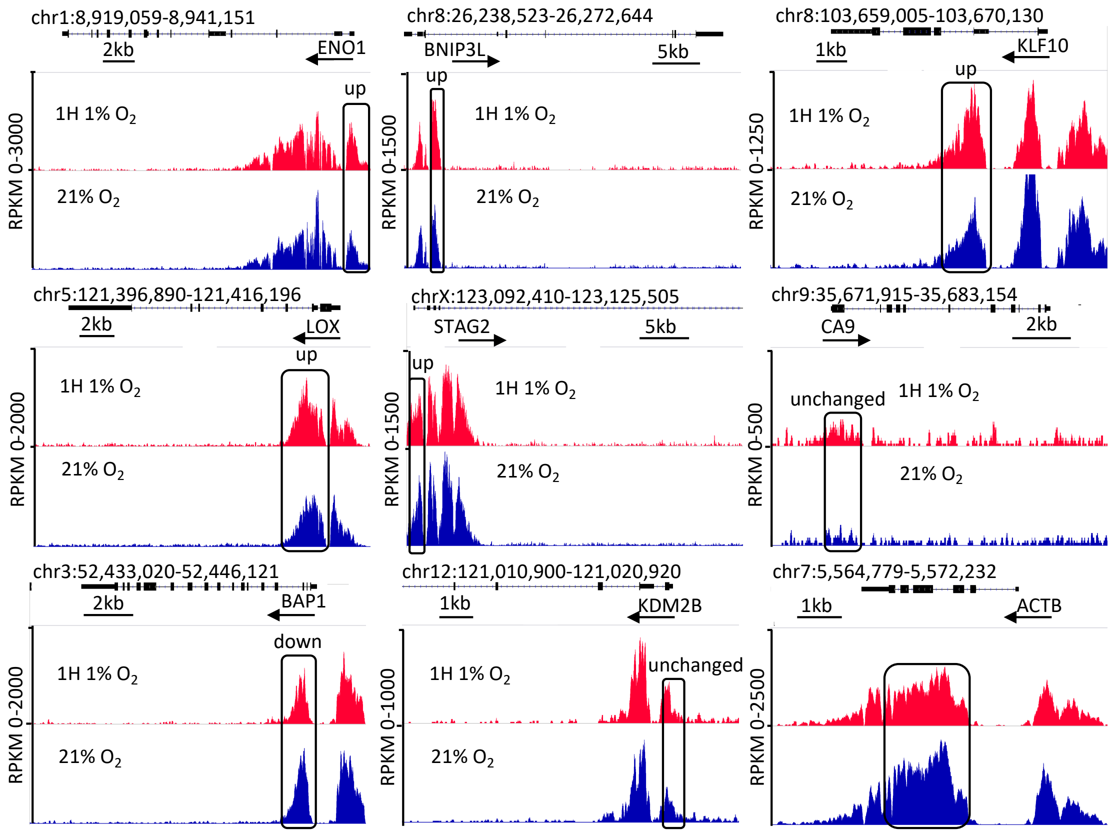
A
chr1:8,919,059-8,941,151
2kb
ENO1
up
1H 1% O2
21% O2
RPKM 0-3000
chr8:26,238,523-26,272,644
BNIP3L
5kb
up
1H 1% O2
21% O2
RPKM 0-1500
chr8:103,659,005-103,670,130
1kb
KLF10
up
1H 1% O2
21% O2
RPKM 0-1250
chr5:121,396,890-121,416,196
2kb
LOX
up
1H 1% O2
21% O2
RPKM 0-2000
chrX:123,092,410-123,125,505
STAG2
5kb
up
1H 1% O2
21% O2
RPKM 0-1500
chr9:35,671,915-35,683,154
CA9
2kb
unchanged
1H 1% O2
21% O2
RPKM 0-500
chr3:52,433,020-52,446,121
2kb
BAP1
down
1H 1% O2
21% O2
RPKM 0-2000
chr12:121,010,900-121,020,920
1kb
KDM2B
unchanged
1H 1% O2
21% O2
RPKM 0-1000
chr7:5,564,779-5,572,232
1kb
ACTB
RPKM 0-2500

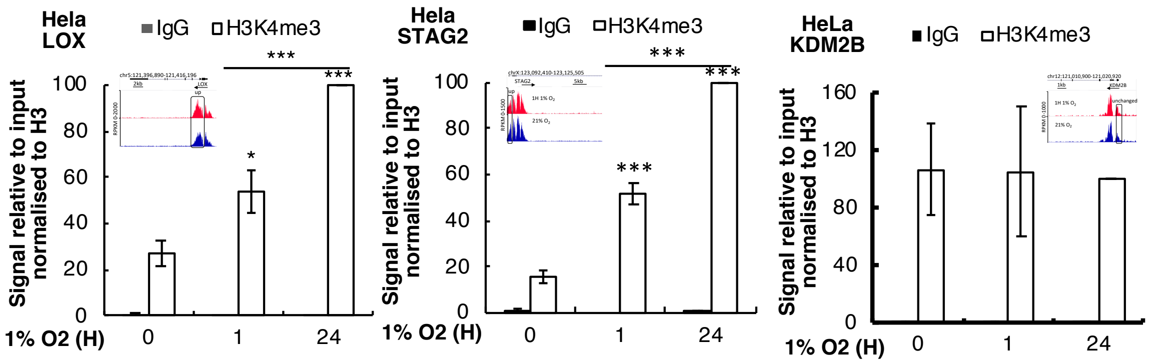
B
Hela LOX
IgG
H3K4me3
Signal relative to input normalised to H3
100
80
60
40
20
0
*
***
***
1% O2 (H)
0
1
24
Hela STAG2
IgG
H3K4me3
Signal relative to input normalised to H3
100
80
60
40
20
0
***
***
***
1% O2 (H)
0
1
24
HeLa KDM2B
IgG
H3K4me3
Signal relative to input normalised to H3
160
120
80
40
0
1% O2 (H)
0
1
24

**Fig. S9.**

**H3K4me3 levels increase in hypoxia at a subset of hypoxia inducible genes.**

Coverage tracks for RPKM normalised H3K4me3 ChIP-Seq read counts in HeLa cells at 21% $O_2$ (normoxia (N) or exposed to 1% $O_2$ for 1 hour (hypoxia (H), at the indicated genes, with hypoxia upregulated peaks outlined (**A**). ChIP-qPCR analysis of H3K4me3 at promoters of the indicated genes in HeLa cells exposed to 1% $O_2$ for the indicated time points (**B**). Graph depicts mean and SEM from a minimum of three independent experiments. Student's t test determined statistical significance. *$p < 0.05$, **$p < 0.01$, ***$p < 0.001$

#### Figure S10

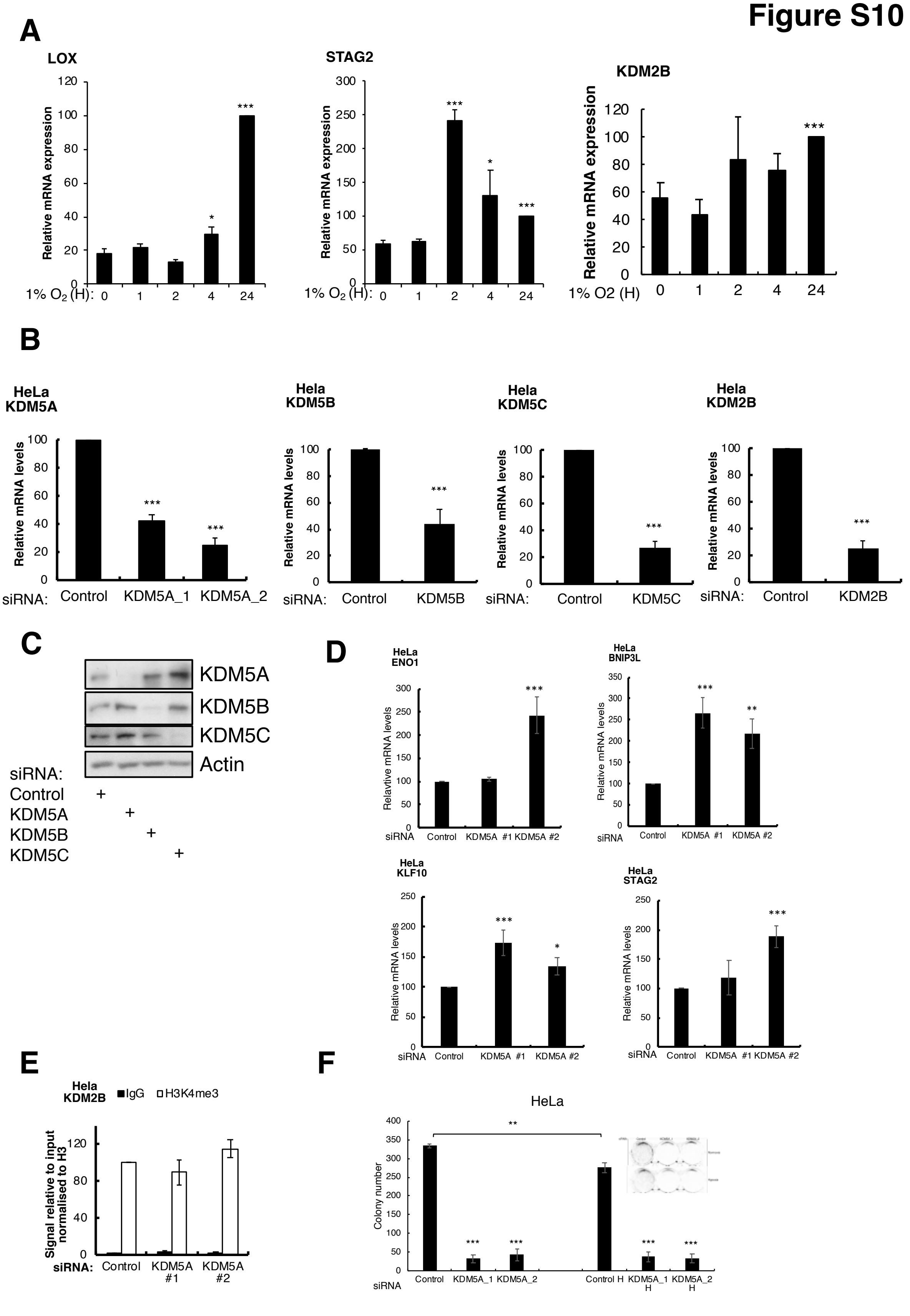

A
LOX
STAG2
KDM2B
Relative mRNA expression
1% O2 (H): 0 1 2 4 24
B
HeLa KDM5A
Hela KDM5B
Hela KDM5C
Hela KDM2B
Relative mRNA levels
siRNA: Control KDM5A_1 KDM5A_2
siRNA: Control KDM5B
siRNA: Control KDM5C
siRNA: Control KDM2B
C
KDM5A
KDM5B
KDM5C
Actin
siRNA:
Control
KDM5A
KDM5B
KDM5C
D
HeLa ENO1
HeLa BNIP3L
HeLa KLF10
HeLa STAG2
Relative mRNA levels
siRNA Control KDM5A #1 KDM5A #2
E
Hela KDM2B
IgG
H3K4me3
Signal relative to input normalised to H3
siRNA: Control KDM5A #1 KDM5A #2
F
HeLa
Colony number
siRNA Control KDM5A_1 KDM5A_2 Control H KDM5A_1 H KDM5A_2 H

**Fig. S10.**
**Validation of siRNA depletion of KDM5 family members.**
qPCR analysis of mRNA expression levels for the indicated genes in HeLa cells exposed to 1% $O_2$ for the indicated time points (A). qPCR analysis of mRNA expression levels for KDM5A, KDM5B, KDM5C and KDM5D in response to siRNA depletion in HeLa cells (**B**). Immunoblot analysis for KDM5 family following siRNA depletion in HeLa cells (**C**). qPCR analysis of mRNA expression levels for the indicated genes in HeLa cells following siRNA-mediated depletion of KDM5A (**D**). ChIP-qPCR analysis for H3K4me3 at the KDM2B promoter following KDM5A depletion (**E**). Graphs depicts mean and SEM from a minimum of three independent experiments. Student's t test determined statistical significance. *$p< 0.05$, **$p< 0.01$, ***$p< 0.001$. Colony formation assay for HeLa cells following siRNA depletion of KDM5A in normoxia and hypoxia. Hypoxia exposure was only conducted for the last 3 days of the 7 days assay (**F**). Graphs depicts mean and SEM from a minimum of three independent experiments. Student's t test determined statistical significance. *$p< 0.05$, **$p< 0.01$, ***$p< 0.001$.

A

### Figure S11

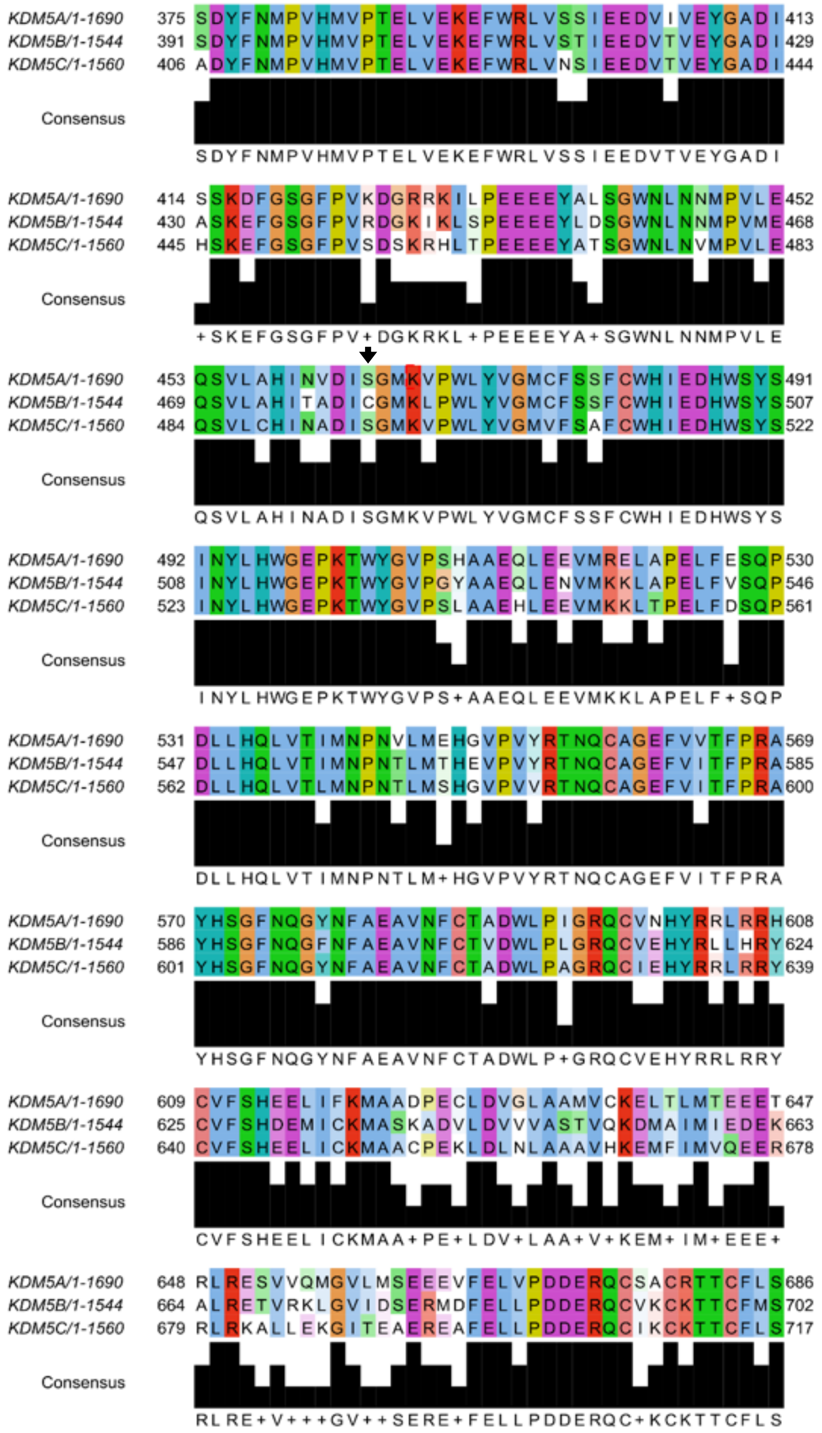

**Fig. S11. Sequence alignment of KDM5A KDM5B and KDM5C**.

Protein sequences from human KDM5A, KDM5B and KDM5C were analyzed for level of homology using Jalview (**A**).

### Figure S12

**A**

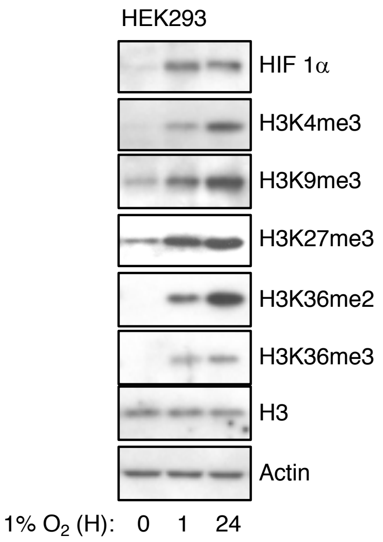


**B**

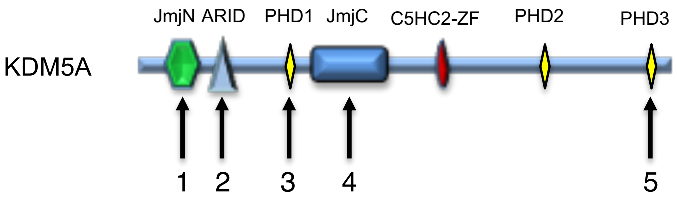


1. T30A, S34A-in KDM5B
2. V109H, as in KDM5B
3. M297L, as in KDM5B
4. S464C, as in KDM5B
5. D1633N, as in KDM5B

**Fig. S12. HEK293 histone mark response to hypoxia and mutation of KDM5A.**

Immunoblot analysis of histone methylation marks in HEK293 following exposure to 1% $O_2$ for the indicated periods of time (**A**). Schematic diagram of KDM5A domain structure and annotation of mutants used in this study (**B**).

### Figure S13

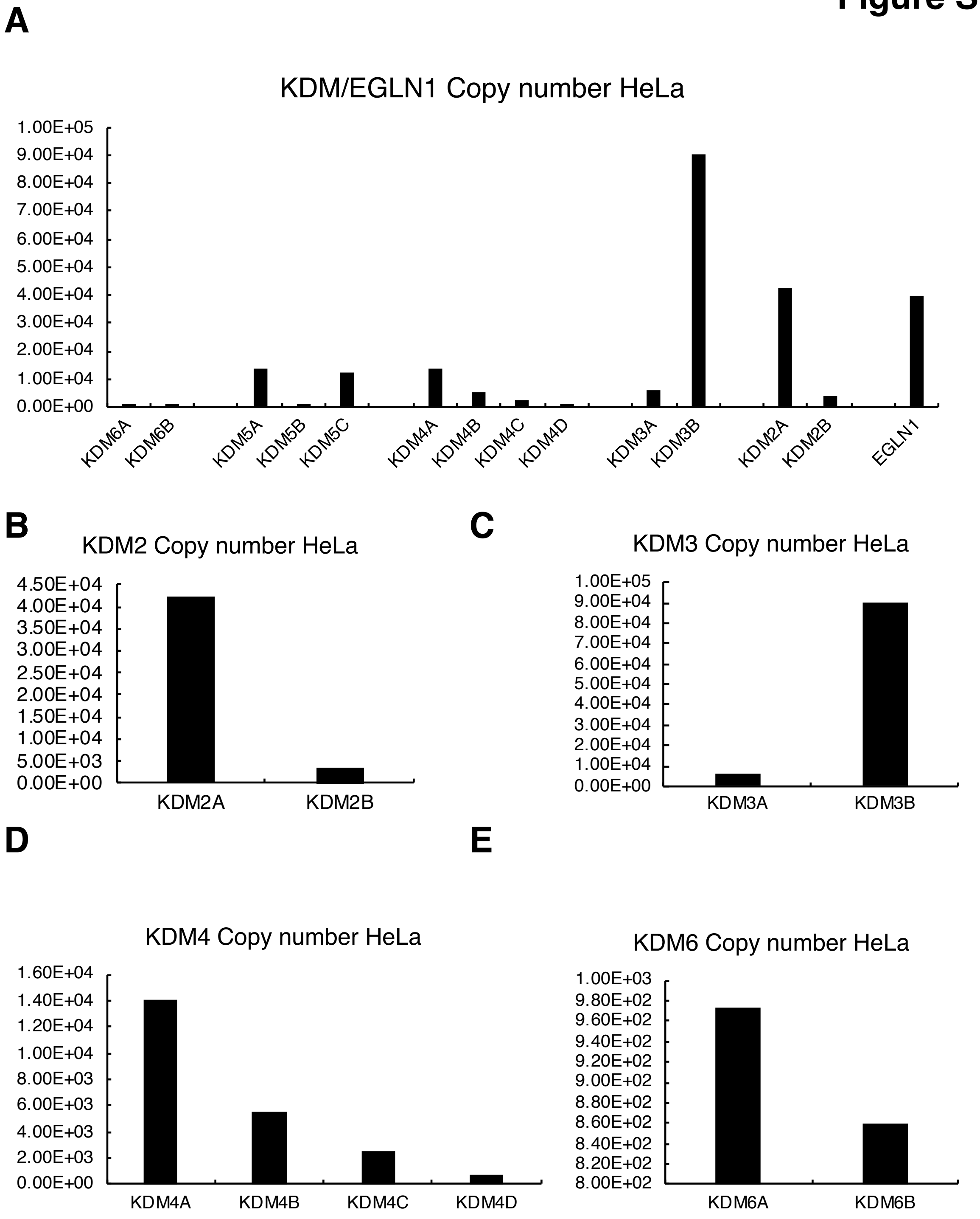

A
KDM/EGLN1 Copy number HeLa
1.00E+05
9.00E+04
8.00E+04
7.00E+04
6.00E+04
5.00E+04
4.00E+04
3.00E+04
2.00E+04
1.00E+04
0.00E+00
KDM6A
KDM6B
KDM5A
KDM5B
KDM5C
KDM4A
KDM4B
KDM4C
KDM4D
KDM3A
KDM3B
KDM2A
KDM2B
EGLN1
B
KDM2 Copy number HeLa
4.50E+04
4.00E+04
3.50E+04
3.00E+04
2.50E+04
2.00E+04
1.50E+04
1.00E+04
5.00E+03
0.00E+00
KDM2A
KDM2B
C
KDM3 Copy number HeLa
1.00E+05
9.00E+04
8.00E+04
7.00E+04
6.00E+04
5.00E+04
4.00E+04
3.00E+04
2.00E+04
1.00E+04
0.00E+00
KDM3A
KDM3B
D
KDM4 Copy number HeLa
1.60E+04
1.40E+04
1.20E+04
1.00E+04
8.00E+03
6.00E+03
4.00E+03
2.00E+03
0.00E+00
KDM4A
KDM4B
KDM4C
KDM4D
E
KDM6 Copy number HeLa
1.00E+03
9.80E+02
9.60E+02
9.40E+02
9.20E+02
9.00E+02
8.80E+02
8.60E+02
8.40E+02
8.20E+02
8.00E+02
KDM6A
KDM6B

**Fig. S13. Quantitative proteomic analysis of KDM family copy number in HeLa cells.**

Copy number analysis for all KDM/EGLN1 members detected and quantified in HeLa cells (**A**). Copy number for KDM2 (**B**), KDM3 (**C**), KDM4 (**D**) and KDM6 (**E**) family members detected and quantified in HeLa cells.

**Data S1. (separate excel file)**

A. HIF 1 validated target genes
B. Conserved hypoxia induced genes
C. Conserved hypoxia repressed genes
D. HeLa hypoxia induced genes
E. HeLa hypoxia repressed genes
F. H3K4me3 normoxic peaks
G. H3K4me3 hypoxic peaks
H. H3K4me3 hypoxia upregulated peaks
I. H3K4me3 hypoxia downregulated peaks
J. H3K36me3 normoxic peaks
L. H3K36me3 hypoxic peaks
L. H3K36me3 hypoxia upregulated peaks
M. H3K36me3 hypoxia upregulated peaks
